# Structural analyses of ‘substrate-pH of activity’ pairing observed in Polysaccharide lyases

**DOI:** 10.1101/2023.02.14.528460

**Authors:** Shubhant Pandey, Bryan W. Berger, Rudresh Acharya

## Abstract

Anionic polysaccharides found in nature are functionally and structurally diverse, and so are the polysaccharide lyases (PLs) which catalyse their degradation. Atomic superposition of various PL folds according to their cleavable substrate structure confirm the occurrence of structural convergence at PL active sites. This suggests that various PL folds have emerged to cleave a particular class of anionic polysaccharide during the course of evolution. While structural and mechanistic similarity of PLs active site have been highlighted in earlier studies, a detailed understanding regarding functional properties of this catalytic convergence remains an open question, especially the role of extrinsic factors such as pH in the context of substrate binding and catalysis. Our earlier structural and functional work on pH directed multi-substrate specificity of Smlt1473 inspired us to regroup PLs according to substrate type to analyse the pH dependence of their catalytic activity. Interestingly, we find that particular groups of substrates are cleaved in a particular pH range (acidic/neutral/basic) irrespective of PLs fold, boosting the idea of functional convergence as well. Based on this observation, we set out to define structurally and computationally the key constituents of an active site among PL families. This study delineates the structural determinants of conserved ‘substrate-pH activity pairing’ within and between PL families.

## Introduction

In our previous study, we have elucidated the structural mechanism for a novel pH directed, multifunctional polysaccharide Lyase (PL) Smlt1473 (PL-5) from *Stenotrophomonas maltophilia* K279a [1]. It specifically and optimally cleaves poly-hyaluronate, poly-glucuronate, and poly-mannuronate at pH 5.0, pH 7.0, and pH 9.0 respectively [2, 3]. The unique pH dependence on catalytic activity enabled us to trap bound poly-mannuronate at non-catalytically active pH 5 and pH 7 in the wild-type Smlt1473 crystal lattice [1]. We furthermore compared the optimum pH for catalytic activity towards a given substrate with that of other monospecific PL families. Like Smlt1473, the PL-8 family of PLs cleaves poly-hyaluronate at acidic pH [4-6], and PL-5 family of PLs cleaves poly-mannuronate at basic pH [7-9].

Motivated by our prior observations, we divided the PLs on the basis of substrate groups as discussed previously [10]. The substrate grouping is based on type of substrate that PLs bind at the [+1] sub-site, and includes- (i) galacturonic acid (GalA), (ii) glucuronic/iduronic acid (GlcA/IdoA), and (iii) guluronic/mannuronic acid (GulA/ManA) (FIGURE 1). In grouping PLs according to substrate type, we have tabulated the information such as substrate group, substrate, PL family, and subfamily, E.C. number, source organism, PDB ID and optimum pH of activity (Table S1). The range of optimum pH for activity for each substrate group establishes the presence of conserved substrate-pH pairing within and among PL families irrespective of variations in structural folds (FIGURE S1). The structural convergence at the active site as a function of substrate has been reviewed previously [10]. However, these prior studies do not explore the associated functional convergence. Thus, based on our prior studies with Smlt1473, we propose pH as a key, functional convergence factor for analysing structurally and computationally the PLs selected from each substrate group.

**FIGURE 1.**
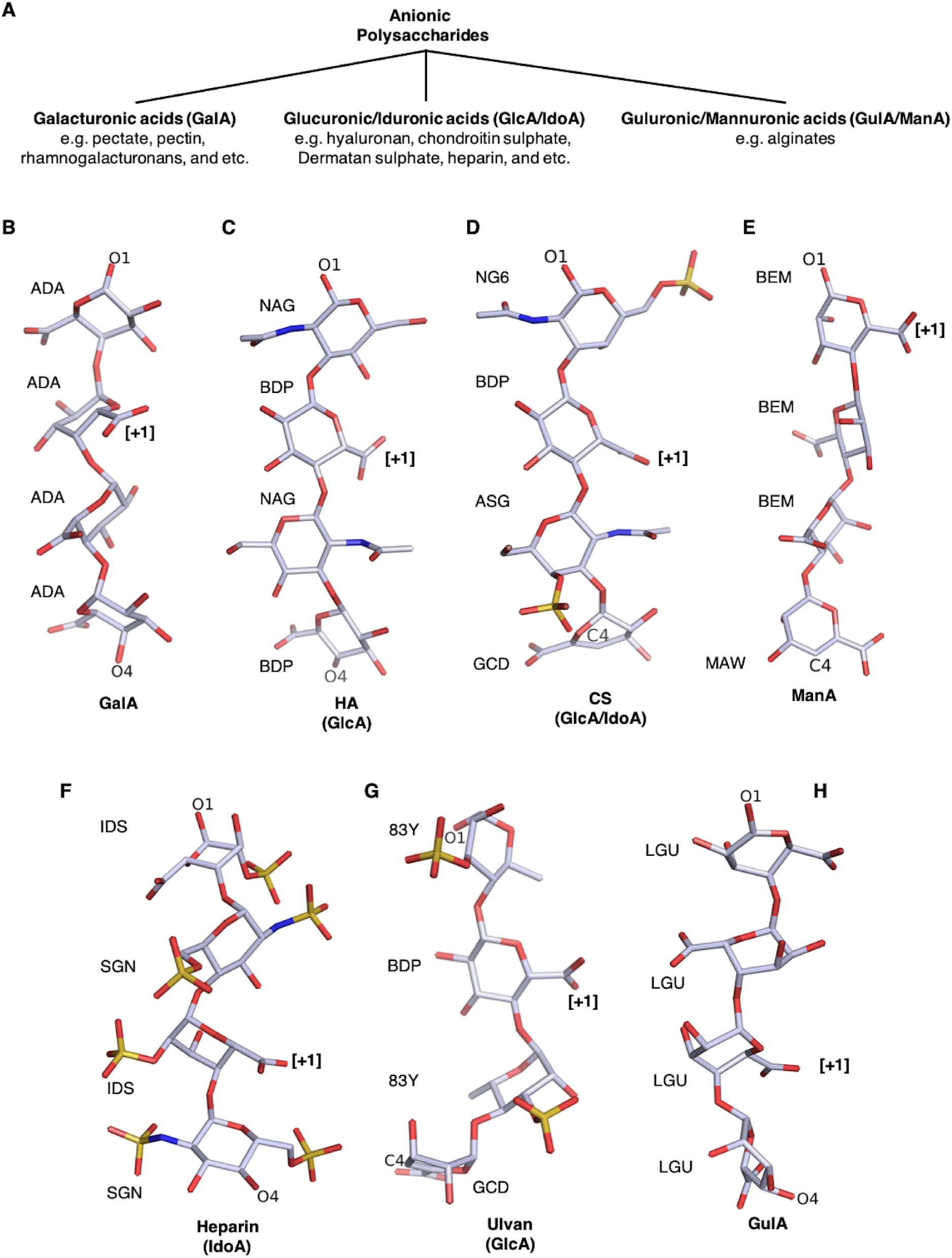
The three main anionic polysaccharide substrate groups are cleaved by PLs. (A) The organization chart depicts the three substrate groups, Galacturonic acid (GalA), Glucuronic/Iduronic acid (GlcA/IdoA), and Guluronic/Mannuronic acid (GulA/ManA). The GalA are modified either as pectate, pectin, or Rhamno-Galacturonan (RG). The GlcA and IdoA are C-5 epimer and similarly are the GulA and ManA. The GalA is a major component of pectic substance found in the cell wall of plants, GlcA/IdoA are the major component of glycans found in the extracellular matrix of higher eukaryotes, GulA/ManA together forms alginates, a major component of seaweed such as brown algae, ulvan are GlcA(s) found in green algae. (B) poly-GalA (pectate), (C) poly-GlcA (HA=Hyaluronic Acid), (D) poly-GlcA (CS=Chondroitin Sulphate), (E) poly-ManA (alginate), (F) poly-IdoA (Heparin), (G) poly-GlcA (ulvan), (H) poly-GulA (alginate). ADA=Alpha-D-galactopyranuronic Acid, NAG= N-acetyl-beta-D-glucosamine, BDP= beta-D-glucopyranuronic acid, NG6=N-acetyl-D-galactosamine 6-sulfate, ASG=N-acetyl-4-O-sulfo-beta-D-galactosamine, GCD=4,5-dehydro-D-glucuronic acid, BEM= beta-D-mannopyranuronic acid, MAW= 4-Deoxy-D-mannopyranuronic acid, IDS= 2-O-sulfo-alpha-L-IDopyranuronic acid, SGN= N, O6-disulfo-Glucosamine, 83Y= 6-deoxy-3-O-sulfo-alpha-L-mannopyranose, LGU= alpha-L-gulopyranuronic acid.

During PLs mediated β-elimination, apart from the first charge neutralization step, the next major and a pre-transition step is C5-H abstraction by a catalytic base from the sugar unit bound at [+1] subsite [10]. As all the catalytic bases in case of PLs are titratable residues (e.g. His, Tyr, Arg, or Lys), their pKa value in the substrate-bound state decides the pH value at which these respective bases get activated. Thus, the pKa value of the residues acting as base defines the base activation pH. At or above this particular pH value these residues will be in a deprotonated form (His, Arg, Lys will become neutral by losing their positive charge, and Tyr will develop negative charge) (FIGURE S2). This is necessary to act as the catalytic base for C5-H abstraction. Based on our work and others, it is evident that the catalytic residues, especially His/Tyr, are involved across the entire pH range of catalytic activity [1-10, 21-28]. Therefore, it is important to analyze structurally and computationally the electrostatic microenvironment of the catalytic bases among selected PLs to determine how differences in microenvironment tune specificity, including both surrounding residues as well as catalytic residues.

If we compare enzymatic activity at different pH values with the idea of an optimum pH observed, the aforementioned base activation pH can be considered as the starting point, the optimum pH for activity can be considered as the intermediate point of highest activity, and outside of this optimum pH enzymatic activity decreases. In other words, the range of pH values where PLs function is known as the pH range for activity. The PL catalytic surface will lose net positively charged surfaces as the pH increases, decreasing the binding interaction energy between negatively charged substrate and PL. At a certain point it will become unfavourable, coinciding with the bounds of the ‘pH range of activity’ (FIGURE S3). However, as prior studies have demonstrated, structurally similar PLs exhibit different functional activities towards substrate and solution conditions, leading to key questions such as (1) what determines the optimal catalytic pH, (2) does optimal catalytic pH coincide with optimal binding energy for a given substrate, and (3) why do different substrates require different ranges of optimal catalytic pH?

Previously in an attempt to address these questions, we had calculated the electrostatic surface map of Smlt1473 wild-type protein structure as a function of pH [1]. We found that at acidic, neutral, and alkaline pH, the respective spread of electropositive surface is maximum, intermediate, and minimum. We anticipate the change in spread of electrostatic surface as we move across the pH range for activity provides a complementary positive surface to PLs for binding anionic polysaccharides, which is the basis for this current study. Our current work suggests the optimum pH corresponds to a pH-directed, electropositive surface where binding is optimal. In order to further clarify the role of pH-directed, charged surfaces in determining the optimum pH range as a function of substrate, we analysed the positive and negative charged amino acids at active and distal sites across a range of PL families and structures. Specifically, we modelled the coulombic interaction network based on our pKa calculations. This allowed us to identify the structural determinants responsible for observed variation in substrate-pH pairing within and among PL families. We also discuss why certain substrates require specific pH ranges for catalysis. Overall, this study will help to create a foundation for PL structure and design which will inspire the development of future research involving engineering substrate specificity into PLs and other carbohydrate-active enzymes where pH plays a significant functional role in catalysis.

## Results and Discussion

### Substrates dictates the PLs activity of pH

As all of the amino acids implicated as the catalytic base for PLs are titratable in nature, the pH at which they can act as base is linked to the magnitude of their pK_a_ values. Thus, amino acids with higher pK_a_ values acts as a base at higher pH and include either Arg (pK_a_=12.50), Lys (pK_a_=10.50), or Tyr (pK_a_=10.00); amino acids acting as base at neutral pH include Lys, Tyr, His (pK_a_=6.50); amino acids acting as base at acidic pH invariably include His. The pK_a_ values shown in bracket are called model or intrinsic pK_a_ values calculated by NMR spectroscopy of terminally capped oligopeptide containing one of the above titratable group **[11, 12]**. At solution pH equal to their pK_a_ values, each amino acid exist in 50% charged, and 50% uncharged state. Thus, at pH values below their pK_a_ value, the amino acids Arg, His, and Lys will remain protonated and carry a net +1-unit positive charge, and cannot acts as base. However, at pH equal or above their pK_a_ values they will be deprotonated and can act as nucleophilic base or proton abstractor. Unlike Arg, His, and Lys, which become neutral after deprotonation at pH above their pK_a_ value, Tyr become negatively charged **(FIGURE S2)**.

At the catalytic site of PLs, the above discussed intrinsic pK_a_ values change in response to solvent accessibility and the electrostatic microenvironment changes drastically in comparison to the fully-solvated, small model oligopeptides [**11-12**]. Moreover, our observation of optimum pH for enzymatic activity for selected PLs belonging to each substrate group is consistent with above discussion **(Table 1, Table S1)**. The PLs cleaving group (i) substrates containing GalA utilizes Arg or Lys as the catalytic base. Notably, the observed optimum pH is below the amino acid’s respective intrinsic pK_a_ values. Moreover, among PLs cleaving a particular substrate at a certain range such as alkaline pH, one can find the optimum pH for activity at much lower or higher pH values. Specifically, in pectate lyases displaying a parallel β-helix fold, in comparison to the lower range of alkaline optimum pHs for activity of PL-3 family member PecB **[13]**, and PL-9 family member BT4170 [**14**], the PL-1 pectate lyases PelC **[15, 16]** and PelA **[17]** show a higher range of alkaline optimum pH values for activity (9.5 and 11.5 respectively). The utilization of lysine as opposed to Arg (PL-1) as catalytic base by PL-3 and PL-9 has been attributed to this observed difference in pH maximum for activity **[13]**. In particular, the low pKa of Lys will lead to its deprotonation at lower pH to act as base, hence the low pH observed for activity **[13]**. However, despite this general trend, we note BsPel (PL-1) has a lower optimum pH of 8.5 as PL-3 and PL-9 but still uses Arg as the catalytic base **[18]**. This observation further validates the microenvironment of the catalytic base is more important than the choice of catalytic base itself in deciding the optimal pH for activity. The other GalA lyase from family PL-2 **[19]** and PL-10 **[20]**, which display structurally dissimilar (α/α)_n_ barrel folds, also utilizes Arg as the catalytic base and like the aforementioned parallel β-helix GalA lyase, utilizes metal ion for substrate binding, and has optimum pH for activity in higher alkaline range as does PelA and PelC **(FIGURE 2)**.

**Table 1.**
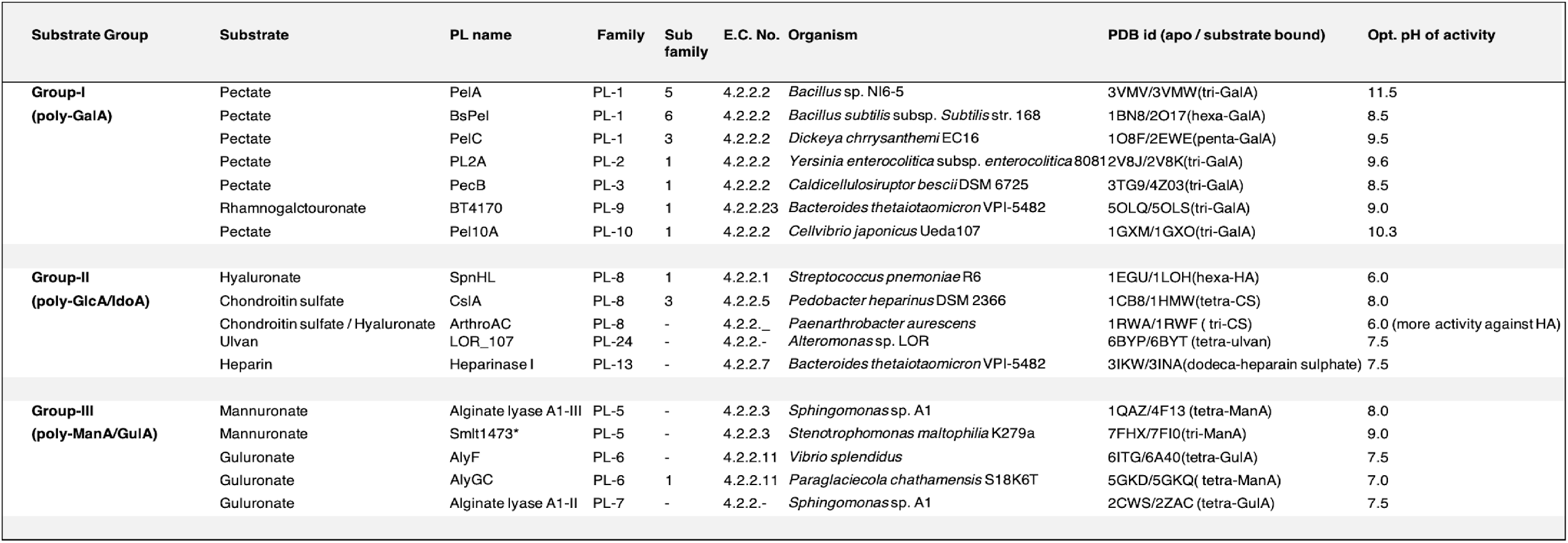
PLs selected for structural and computational analyses.

**FIGURE 2.**
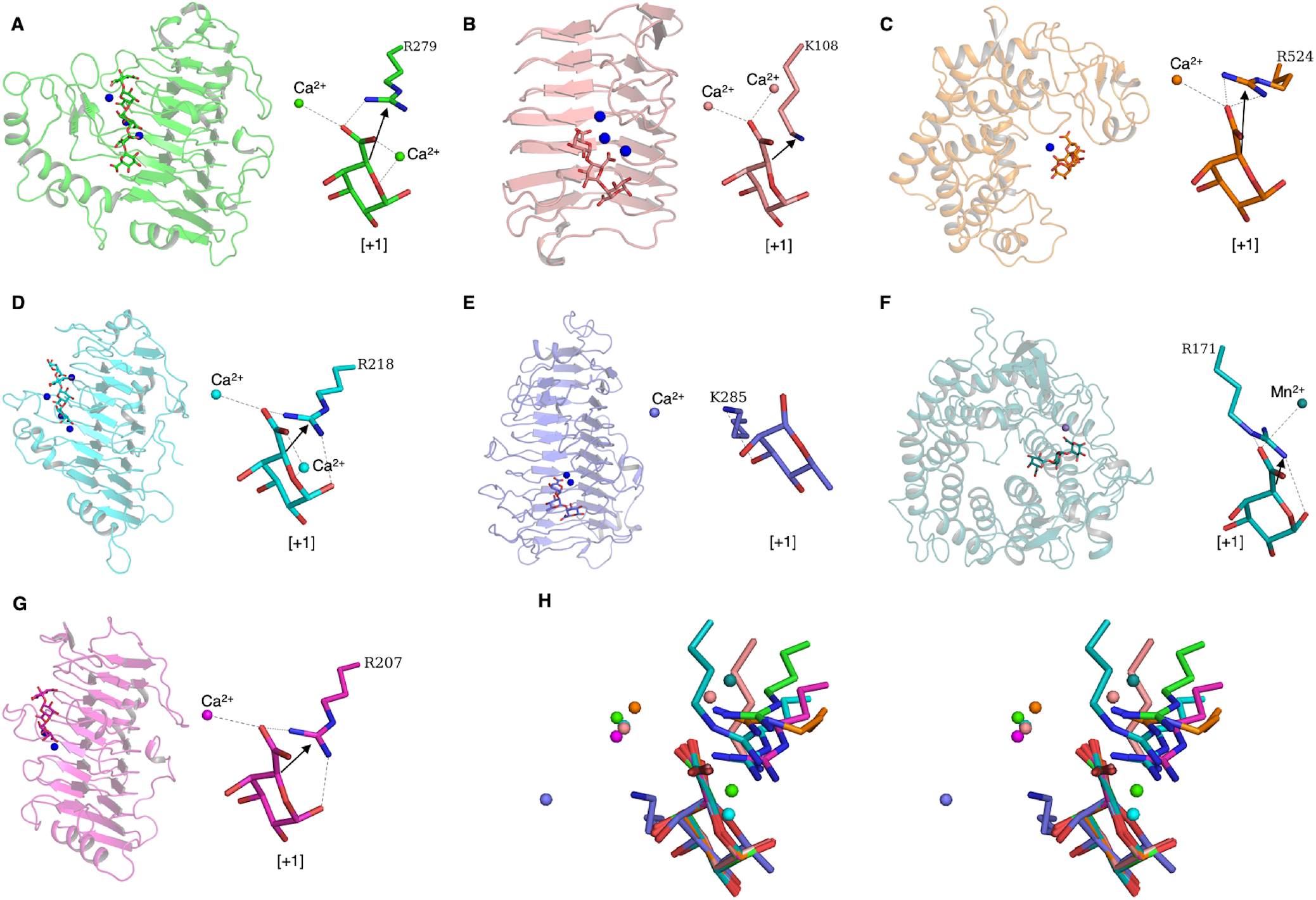
The Poly-GalA lyase from various PL families. The cartoon representation of the PL fold and stick representation of the active site at [+1] subsite of- **(A)** BsPel from *Bacillus subtilis* subsp. *Subtilis* str. 168, PL-1, subfamily-6, PDB id: 1BN8/2O17; **(B)** PecB from *Caldicellulosiruptor bescii* DSM 6725, PL-3, subfamily-1, PDB id: 3T9G/4Z03; **(C)** Pel10A Pel10A from *Cellvibrio japonicus* Ueda107, PL-10, subfamily-1, PDB id: 1GXM/1GXO, **(D)** PelC from *Dickeya chrysanthemi* EC16, PL-1, subfamily-3, PDB id: 1O8F/2EWE; **(E)** BT4170 from *Bacteroides thetaiotaomicron* VPI-5482, PL-9, subfamily-1, PDB id: 5OLQ/5OLS; **(F)** PL2A PL2A from *Yersinia enterocolitica* subsp. *enterocolitica* 8081 PL-2, subfamily-1, PDB id: 2V8J/2V8K; **(G)** PelA from Bacillus sp. N16-5, PL-1, subfamily-5, PDB id: 3VMV/3VMW; **(H)** stereographic image of the superposing substrate (+1 subsite) from all poly-GalA lyase (A-G) for visualizing structural convergence at the active site.

Among PLs cleaving group (ii) substrate (GlcA/IdoA), the PL-8 family of PLs displaying (α/α)_n_ barrel and anti-parallel β-sheets cleave hyaluronic acid at acidic pH (SpnHL, ArthroAC, ScPL8Hyal) but cleave chondroitin sulphate (CslA) at alkaline pH. Like PL-5 enzymes, the PL-8 family of PLs are known to utilize His and Tyr as catalytic base and acid respectively **[4-6, 21-23]**. However, the only difference being that PL-8 His and Tyr are in *syn* position compare to *anti* of PL-5. Interestingly, another chondroitin sulphate lyase from family PL-6, which display parallel β-helix like pectate lyase, use metal ions for substrate binding, Arg as catalytic base, and thus utilize alkaline pH for enzymatic activity **[24, 25] (Table 1, Table S1)**. The heparin lyase Heparinase-I (PL-24) cleaves the most negatively charged anionic polysaccharide heparin sulphate near neutral pH of 7.5 **[26]**. It displays β-jellyroll fold, and utilizes His/Tyr system similar to Smlt1473. The last selected PL from this group include ulvan lyase LOR_107 (PL-13), cleaves sulphated GlcA substrate at pH-7.5, and displays a less characterized β-propeller fold. It also contains structurally similar His/Tyr system as PL-8 enzymes, but His has been implicated to act as both acid and base **[27] (FIGURE 3)**.

**FIGURE 3.**
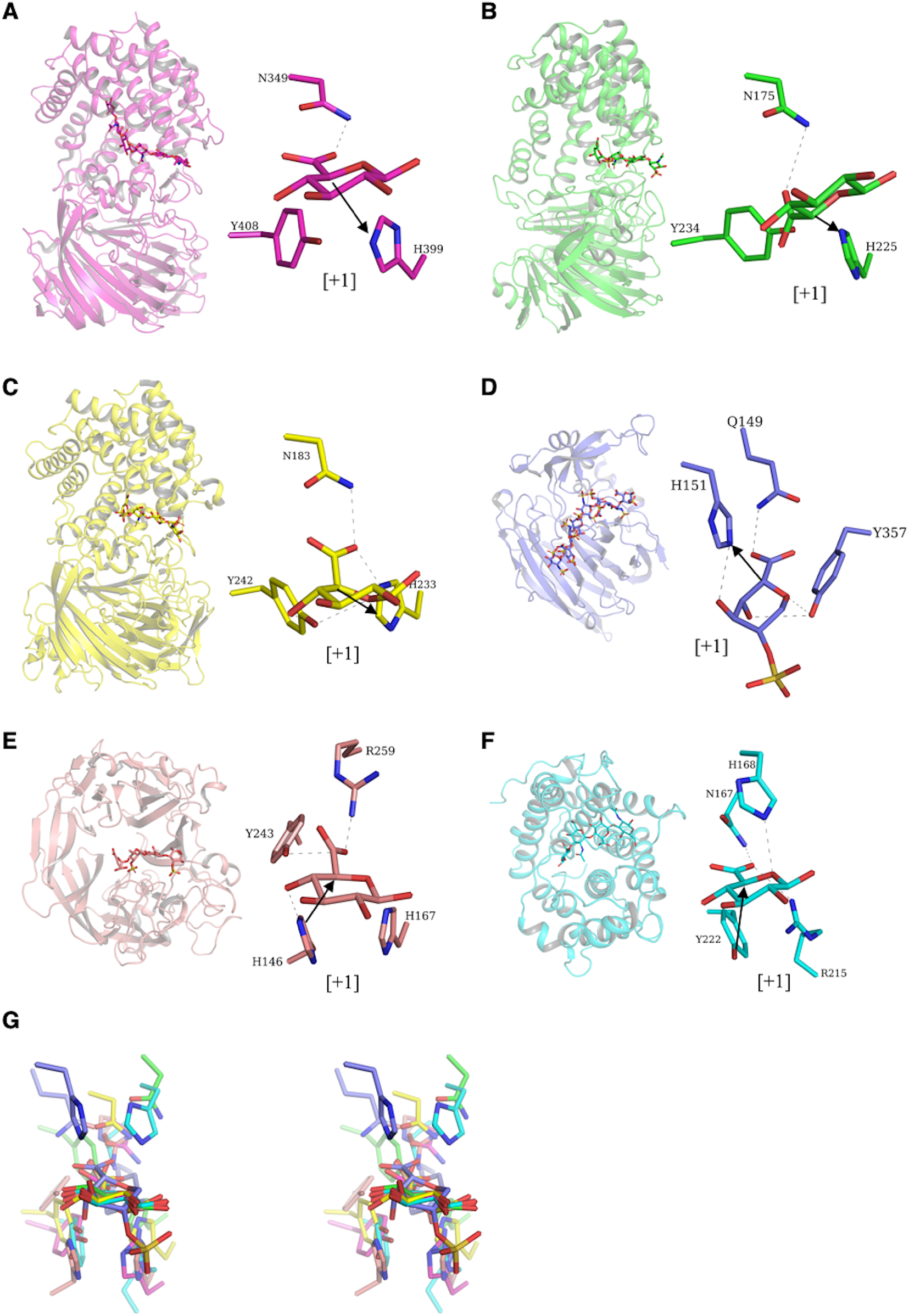
The Poly-GlcA/IdoA lyase from various PL families. The cartoon representation of the PL fold and stick representation of the active site at [+1] subsite of- **(A**) SpnHL from *Streptococcus pneumoniae* R6, PL8, subfamily-1, PDB id: 1EGU (apo)/1LOH (hexa-HA); **(B)** CslA from *Pedobacter heparinus* DSM 2366, PL8, subfamily-3, PDB id: 1CB8 (apo)/1HMW (tetra-CS); **(C)** ArthroAC from *Paenarthrobacter aurescens*, PL8, PDB id: 1RWA (apo)/1RWF (iduronated CS); **(D)** Heparinase I from *Bacteroides thetaiotaomicron* VPI-5482, PL13, PDB id: 3IKW (apo)/3INA (Heparin); **(E)** LOR_107 from *Alteromonas sp*. LOR, PL-24, PDB id: 6BYP (apo)/ 6BYT (ulvan); **(F)** Smlt1473 from *Stenotrophomonas maltophilia* K279a, PL5, PDB id: 7FHX (apo pH-5)/7FHX (docked HA); **(G)** stereographic image of superpose substrate (+1 subsite) from all poly-GlcA/IdoA lyase (A-F) for visualizing structural convergence at active site.

Among PLs cleaving group (iii) substrate (ManA/GulA), the PLs selected for ManA include only well characterized PL-5 enzymes, whereas for GulA we have included PL-6 and PL-7 enzymes. The PL-5 family member Smlt1473 alginate lyase A1-III [(α/α)_n_ barrel] and PL-7 alginate lyase A1-II (β-sandwich) are structurally diverse but have a similar His/Tyr catalytic system **[1, 7-9, 28]**. The PL-6 AlyF displays a parallel β-helix as pectate lyase, and similarly utilizes Arg as the catalytic base but does not use metal ions for ligand binding **[29]**. Again, the optimum pH for ManA lyase is in the alkaline pH range, and in spite of structural divergence PL-6 (parallel β- helix) and PL-7 (β-sandwich) utilize the same optimum pH of 7.5 **(FIGURE 4)**.

**FIGURE 4.**
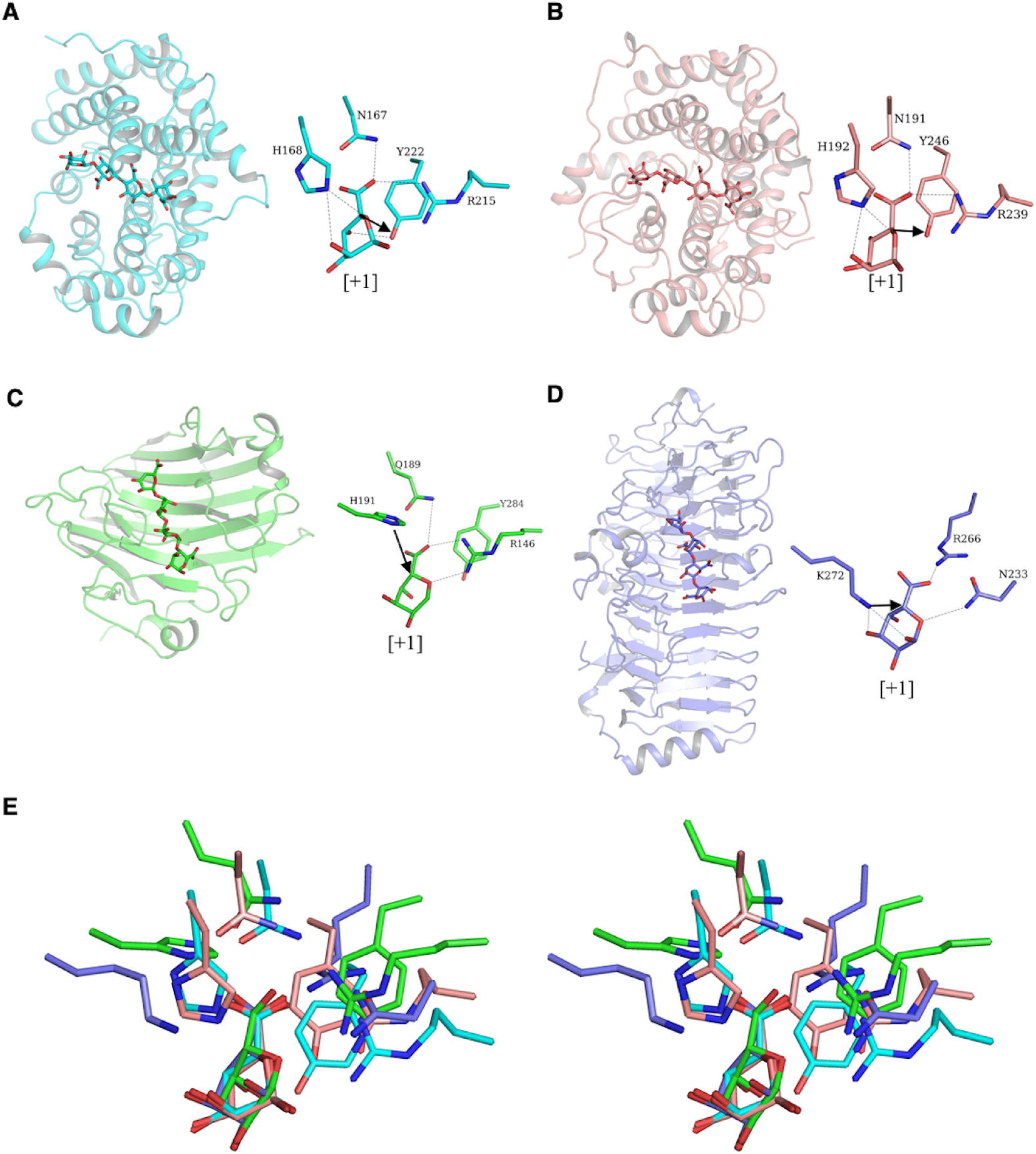
The Poly-GulA/ManA lyase from various PL families. The cartoon representation of the PL fold and stick representation of the active site at [+1] subsite of- (A) Smlt1473 from *Stenotrophomonas maltophilia* K279a, PL5, PDB id: 7FHZ (apo at pH-9)/7F10 (ManA bound at pH-5.0), 7FI1(ManA at pH-7.0); (B) Alginate lyase A1-III from *Sphingomonas* sp. A1, PL-5, PDB id: 1QAZ (apo)/4F13 (ManA); (C) Alginate lyase A1-II from *Sphingomonas* sp. A1, PL-7, PDB id: 2CWS (apo)/2ZAA (ManA); (D) AlyF from *Vibrio splendidus*, PL-6, PDB id: 6ITG (apo), 6A40 (GulA); (E) stereographic image of superpose substrate (+1 subsite) from all poly-ManA/GulA lyase (A-D) for visualizing structural convergence at active site.

We found from the above structural analysis that the PLs with parallel β-helix fold cleave substrates from each group such as pectate (GalA), chondroitin sulphate (GlcA/IdoA), and poly-GulA (GulA/ManA,) but retain the Lys/Arg catalytic system. Except for AlyF (poly-GulA lyase, optimum pH=7.5), they utilize metal ions for substrate binding and charge neutralization, and maintain an alkaline range of pH for optimal activity. On the other hand, the PLs displaying (α/α)_n_ barrel fold utilize different catalytic mechanism for GalA (Lys/Arg system, metal ion, alkaline pH for optimal activity) in comparison to GlcA/IdoA, and GulA/ManA (His/Tyr system, substrate specific pH for optimal activity). Therefore, we observed that PLs displaying (α/α)_n_ barrel fold show high divergence at the active site and thus maintain various ranges of substrate-specific optimum pH values **(Table 1)**. This is in contrast to PLs displaying parallel β-helix folds, which show high convergence at the active site, but still maintain various substrate-specific optimum pH values by subtle arrangements of flanking residues **(Table 1)**. The other selected PLs, which contain folds such as β-propeller (LOR_107, a PL23 ulvan lyase), β-sandwich (alginate lyase A1-II, a PL-7 poly-GulA lyse), or β- jellyroll (Heparinase-I, a PL-13 heparin sulphate lyase), utilize a His/Tyr system and show neutral and near neutral range of optimum pH for activity. Finally, we conclude that structural convergence may not entirely coincide with pH dependent activity-based functional convergence. In contrast, we find that, irrespective of convergence and divergence in their fold and catalytic geometry, the PLs have evolved to structurally and mechanistically adjust themselves to maintain a strict substrate-pH pairing. The pK_a_ of PL’s catalytic base is determined by the electrostatic interactions available at active site after substrate binding. In order to understand what affects the pK_a_ values of PL catalytic base residues, we have performed structural modelling and computational pK_a_ analyses of selected PLs (See Materials and Methods). A brief primer on general theories (Born effect, H-bonding, coulombic interactions) regarding pK_a_ perturbation is detailed in the supplement (Note S1).

### PLs binding GalA at [+1] subsite, displaying parallel β-helix fold

The most common feature among these PLs is the presence of metal ions (Ca^2+^ and Mn^2+^) mainly at [-1, +1, +2] subsite **(FIGURE 2, 5)**. Metal ions most often play a critical role in substrate binding, positioning, and charge neutralization step of the β-elimination reaction mechanism. The other common feature of these PLs is utilization of basic amino acids Arg or Lys as the catalytic base, and utilization of alkaline pH for catalytic turnover **[13-20]**. The in-depth structural description of fold architectures among these PLs are provided in supplementary **(Note S2, FIGURE S3A-B)**. As discussed previously, the PLs cleaving GalA containing substrates occupy different optimum pH positions in the alkaline pH range. The PLs (PL-3, and PL-9) occupying a lower alkaline pH range (8-9) generally use Lys as catalytic base, and PLs (PL-1, PL-2, PL-10) occupying higher alkaline pH range (9.5-11.5) use Arg as catalytic base **[13]**. However, in our analysis we found a poly-GalA lyase BsPel which utilize Arg as catalytic base, utilizes a lower alkaline pH range of optimum activity (8.5) as PL-3 and PL-9 **[18]**. Thus, PL-1 family of pectate lyases provide a first opportunity to discern the mechanism of the wide optimum pH variation (8.5-11.5) within a PL family. We performed sequence and structural comparisons and identified some key similarity and differences. PelA (sf-5) has minimal T3 insertion followed by PelC (sf-3), whereas BsPel (sf-6) has significant T3 insertion **(FIGURE S3D-E, H-I)**. The structural alignment in terms of r.m.s.d has the following order: [PelC-PelA] > [BsPel-PelA] > [PelC-BsPel]. The PelA being intermediate subfamily-5 superimpose well with both PelC and BsPel, but there is significant r.m.s.d between PelC and BsPel **(FIGURE S3C-S3E)**. A similar trend is followed for substrate-bound structures as well **(FIGURE S1G-S1I)**, however removal of T3 insertion from both PelC and BsPel results in significantly improved r.m.s.d **(FIGURE S3F)**. Our further analyses of active site constituent amino acids among these PLs reveal the role of T3 insertion in shaping the electrostatic microenvironment by adding Ca^2+^ binding amino acid residues. In contrast to PelA, the T3 insertion contribute pair of amino acids for both PelC (D160, D162) and BsPel (D173, N180) which help them bind Ca^2+^ ion at [+1] subsite **(FIGURE S4A-C)**.

**FIGURE 5.**
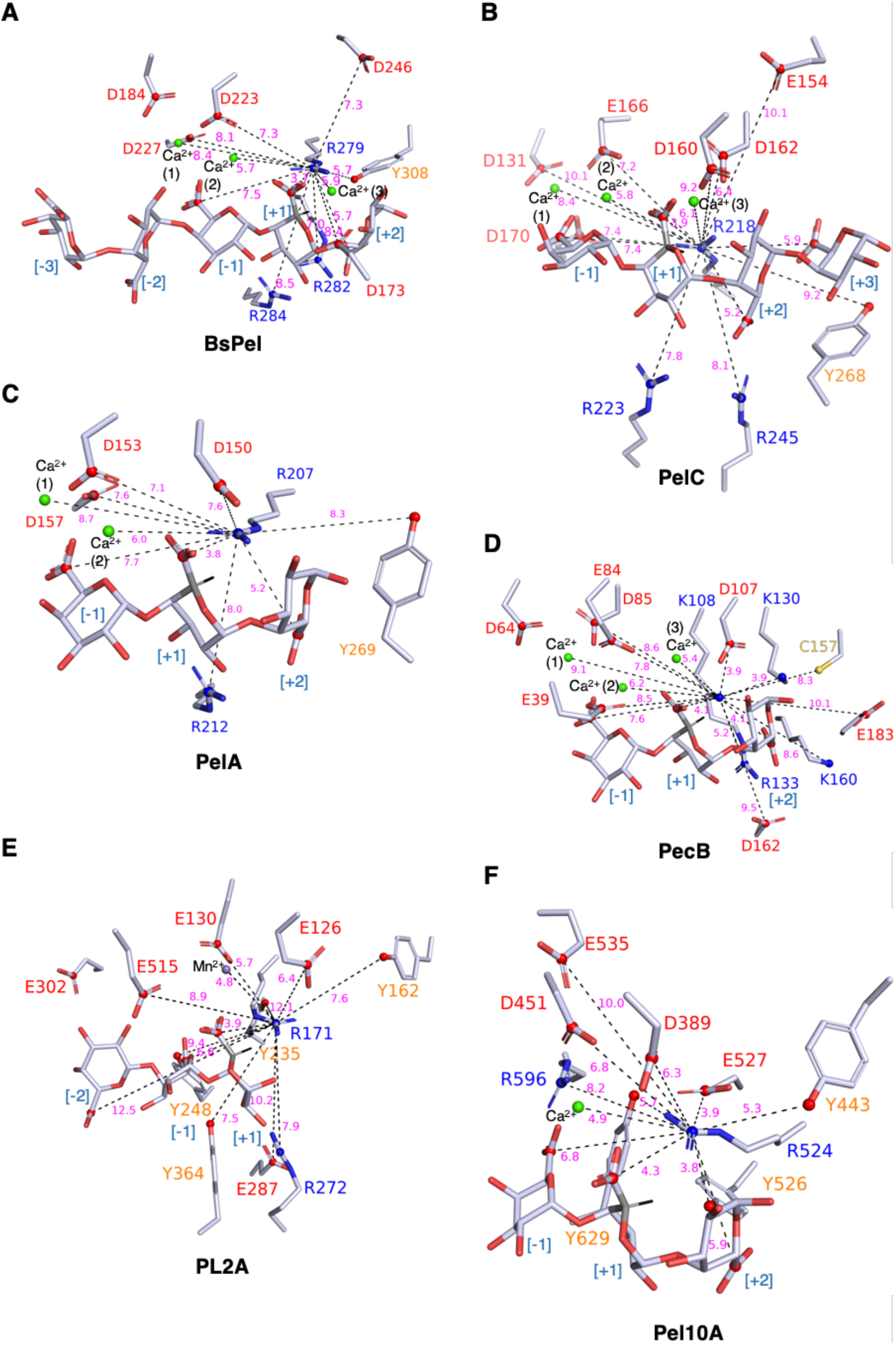
Electrostatic microenvironment around the catalytic base of selected Poly-GalA lyases. (A) BsPel complexed with penta-GalA [-3 to +2], and three Ca^2+^ ion [-1 to +1], catalytic base: R279 [+1], (B) PelC complexed with tetra-GalA [-1 to +3], and three Ca^2+^ ion [-1 to +1], catalytic base: R218 [+1], (C) PelA complexed with tri-GalA [-1 to +2], two Ca^2+^ ion [-1], catalytic base: R207, (D) PecB complexed with tri-GalA [-1 to +2], three Ca^2+^ ion [-1 to +1], catalytic base: K108, (E) PL2A complexed with tri-GalA [-2 to +1], Mn^2+^ ion [+1], catalytic base: R171, (F) Pel10A complexed with tri-GalA [-1 to +2], one Ca^2+^ ion [-1], catalytic base: R524. The coulombic interactions centred at the catalytic base are shown as dashed lines (black) labelled with distance (magenta). The basic, acidic, and aromatic amino acids constituting the electrostatic microenvironment are represented as stick model (C=grey, O=red, and N=blue) and respectively labelled with blue, red and orange colours. The substrates are also shown as the same stick model and labelled with PL’s active site surface subsites, negative (left, entry site) to positive (right, exit site). The blue and red non-bonded spheres represent respectively the positive charge centre of basic amino acids (pK_a_ lowering), and the negative charge centre of the carboxylic group of acidic amino acids, and substrates (pk_a_ raising). The cleavable glycosidic oxygen lies in between [-1, +1] subsite, the catalytically important substrate’s C5 atom at [+1] subsite is shown in grey colour with bonded labile H atom represented as a black line.

In case of BsPel, the catalytic base R279 in the apo state face pKa lowering determinants which include positive coulombic interactions from a bound Ca2+ ion, and R284 at [+1] subsite. The determinants of increased pKa include negative coulombic interactions from D184, D223, D227 at [-1] subsite, and D173, D246, Y308 at [+2] subsite **(Table S2)**. The BsPel in the substrate-bound state coordinate two more calcium ions at [-1] subsite; the pKa-lowering determinant R284 and Ca2+ ions are the same as the apo state. The pKa raising determinants apart from D173, D223, D227, D246, and Y308 include additional negative columbic interaction from the substrate carboxylic groups at the [-1, +1, +2] subsites. The negatively charged residue D184 does not interact with R279 in the substrate bound state, as it is shielded by two extra Ca2+ ions binding at [-1] subsite **(Table S2, FIGURE 5A)**.

Similar to BsPel, the PelC catalytic base R218 at [+1] subsite in apo state also faces one Ca^2+^ ion, and a basic amino acid R245 at [+1] subsite. The pKa increasing determinants include D131, E166, D170 at [-1] subsite, D162 at [+2] subsite, and Y268 at [+3] subsite. In the bound state, the pKa increasing determinants include two additional bound Ca^2+^ ions at [-1] subsite, and R245 **(Table S2)**. In contrast to BsPel, the PelC in the bound state interacts with extra negatively charged residues apart from D131, D162, E166, D170, and Y268. The additional negatively charged residues include E154, and D160 at [+2] subsite. Moreover, there is a greater number of substrate-interacting carboxylic groups for PelC at [-1, +1, +2, +3] subsite in comparison to BsPel **(Table S2, FIGURE 5B)**.

As discussed above **(FIGURE S4A-C)**, the PelA catalytic base R207 at [+1] subsite in apo state does not have a conserved Ca^2+^ binding site. The only pKa lowering residue is R212, whereas pKa increasing determinants include D153, D157 at [-1] subsite, D150 at [+1] subsite, and Y269 at [+3] subsite **(Table S2)**. In the bound state, the pKa increasing determinants include two Ca^2+^ ion, and residue R212. The pKa raising determinants include the same amino acid residues as the apo state, namely D153, D157, D150, and Y269. The contributions from bound substrate include carboxylic groups from [-1, +1, +2] subsite **(Table S2, FIGURE 5C)**. However, we can anticipate more electronegative contribution from substrate occupied at other positive subsites (not modelled in the relaxed structure) as the original PDB deposited structure where co-crystallized product is complexed without Ca^2+^ ion, the substrate occupies [+2, +3, +4] subsites **[17]**.

From the above results, it is clear that the structural differences at the active site due to sequence insertion **(FIGURE S3A-B)** can led to considerable changes which affect the substrate binding property of active site within the same family of PL. We observed that Ca^2+^ ions are mainly bound at [-1, +1] subsite in all selected PL-1 family members of PLs. In case of BsPel, they are able to shield the significant negative charge contribution from the acidic amino acids, and substrate carboxylic group from negative subsites [-]. In contrast, PelA and PelC, where substrate mostly occupy [+] subsites, the bound Ca^2+^ ions are not able to shield the negatively charged contribution to the catalytic base. Moreover, like BsPel and PelC, the PelA does not bound Ca^2+^ ion at [+1] subsite, which further led to increase in negative charge contribution to its catalytic base. Thus, it is evident why PelA has higher position in alkaline range of activity followed by PelC, and BsPel based on structural differences at the active site due to sequence insertion and Ca^2+^ ion binding.

The other family of pectate lyases which show a parallel β-helix include, family PL-3 (PecB), and PL-9 (BT4170), which as discussed above utilize the Lys residue as catalytic base whose solution pKa is usually lower than that of Arg. The PecB in apo state binds Ca^2+^ ion at the [+1] subsite, the other pKa lowering determinant for catalytic base K108 include R133 at [+2] subsite, and K130, K160 at [+3] subsite. The pKa raising determinants include, E39, D64, E84, and D85 at [-1] subsite, and D107 at [+2] subsite **(Table S3)**. In the bound state, the PecB binds two more Ca^2+^ ions at [-1] subsite, and retains the same basic residues as pKa raising determinant, R133, K130, and K160. While the number of pKa raising determinants have increased, apart from E39, E84, D85, and D107, the new contributors include residues at and toward [+3] subsite, C157, D162, and E183. The residue D64 does not contribute in the bound state as the Ca^2+^ ion at [-1] subsite shield the possible interaction **(Table S3, FIGURE 5D)**; a similar occurrence has been explained above in case of BsPel **(FIGURE 5A)**. However, the increase in the number of negative charge interactions upon substrate binding is similar to that of PelC. Interestingly, in both cases, these negatively charged substrates are at [+] subsites indicating their possible role in product release. The other PecB pKa raising contribution come from all carboxylic groups of bound substrates at [-1, +1, +2] subsites. As we do not have GalA substrate modelled in case of BT4170 structure at [+1] subsite, we have not performed in depth structural and computational work for comparative analyses.

### PLs binding GalA at [+1] subsite, displaying (α/α)_n_ barrel fold

The last structurally and biochemically characterized poly-GalA lyase selected for analyses include the PLs of family PL-2 (PL2A) and PL-10 (Pel10A), which display (α/α)_n_ barrel fold. Like PL-1, they (PL2A and Pel10A) utilize Arg as the catalytic base, possess a high optimum pH for activity, and depend upon divalent cations for their catalytic activity. The Pel10A, like PL-1 pectate lyase, utilizes Ca^2+^ ion, whereas PL2A utilizes Mn^2+^ ion for charge neutralization. The PL2A in apo state has one Ca^2+^ ion at [-1] subsite as pKa lowering determinants. In contrast, the pKa raising determinants for catalytic base R524 include, H-bond interaction from E527, and Y629 at [+1], and [-1] subsite respectively. The pKa raising determinants involved in electrostatic interactions include, D389, E527, Y526 at [+1] subsite, D451, Y629 at [-1] subsite and, Y443 at [+2] subsite **(Table S3)**. In bound state, apart from Ca^2+^ ion, R596 at [-1] subsite is another pKa lowering determinant. The pKa raising determinants are the same as the apo state, D389, E527, Y526, D451, Y629, and Y443. However, the substrate binding results in one more pKa raising coulombic interaction from E535 oriented towards the [-1] subsite **(Table S3, FIGURE 5E)**.

The Pel10A catalytic base R171 in the apo state has only bound Mn^2+^ ion as pKa lowering determinant; the pKa raising determinant includes E302 toward [-2] subsite, E515, Y235, Y248, Y364 at [-1] subsite, and E130 at [+1] subsite. In the bound state, Mn^2+^ is along with R272 at [+1] subsite as a pKa lowering determinant. The pKa raising determinants apart from E130, E515, Y248, and Y364 include new contribution from E287 at [+1] subsite, and E126, Y162 at [+2] subsite **(Table S3)**. In the bound state, E302, and Y235 do not contribute in raising catalytic base pKa because the rotameric transition of catalytic base toward [+] subsite preclude coulombic interaction from [-] subsites after substrate binding. Similarly, the rotameric transition lead to gain of new interactions toward [+] subsites, E126, E287, and Y162. The carboxylic group from [-1, +1] subsite are other contributors in raising catalytic base pKa **(Table S3, FIGURE 5F)**.

The major difference observed between the poly-GalA lyase of two fundamentally different folds is that PLs displaying parallel β-helix fold coordinate additional metal ions toward [-] subsites after substrate binding compare to PLs displaying (α/α)_n_ barrel fold, which in apo as well as bound state bind only one metal ion at [-1 or +1] subsite. The other major difference is the composition of amino acids at the active site; the PLs displaying parallel β-helix fold have more Asp (D) and less Glu (E) residues as pKa raising determinants. In contrast, the PLs displaying (α/α)_n_ barrel fold has mainly Glu (E), and Tyr (Y) residue as pKa raising determinants. The presence of more Tyr residue is one of the characteristics of the (α/α)_n_ barrel fold, which is apparent in our subsequent analysis of the poly-GlcA/IdoA, and poly-ManA lyase detailed below.

### PLs binding GlcA at [+1] subsite, displaying (α/α)_n_ barrel + antiparallel β sheet fold

The poly-GlcA(s) such as hyaluronic acid (HA), chondroitin sulphate (CS), and ulvans are structurally variable due to their polymeric structure and also due to various degree of substitution replacing sugar OH groups. These substitutions provide each class of sugar distinct chemical properties such as varying negative charge density. As discussed earlier, the aforementioned sugars not only differ in their structures, but also have a very specific range of pH values where PLs will cleave. HA invariably is cleaved at acidic pH, CS based on its composition can have either acidic or alkaline optimum pH, and ulvan is cleaved at near neutral pH. Firstly, we will discuss PL-8 family of poly-GlcA lyase, where we have chosen a HA specific lyase (SpnHL, opt. pH=6.0), CS specific lyase (CslA, opt. pH=8.0), and bi-functional lyase which cleaves both HA and CS (ArthroAC, optimum pH-6.0, more active against HA).

SpnHL in apo state has two pKa lowering determinants for catalytic base H399 at [+1] subsite: R462 at [+1] subsite, and R336 at [+2] subsite. The pKa raising determinants include H-bond interaction and coulombic interaction from E577 at [+1] subsite. Other such coulombic interaction is contributed from D398 at [+1] subsite and E388 at [+2] subsite **(Table S4a)**. In the HA bound state, the pKa lowering determinants are the same as the apo state, R336, and R462. However, the pKa raising determinants includes H-bond interaction with sugar ring oxygen atom at [+1] subsite, and single coulombic interaction from E577 **(Table S4a, FIGURE 6A)**.

**FIGURE 6.**
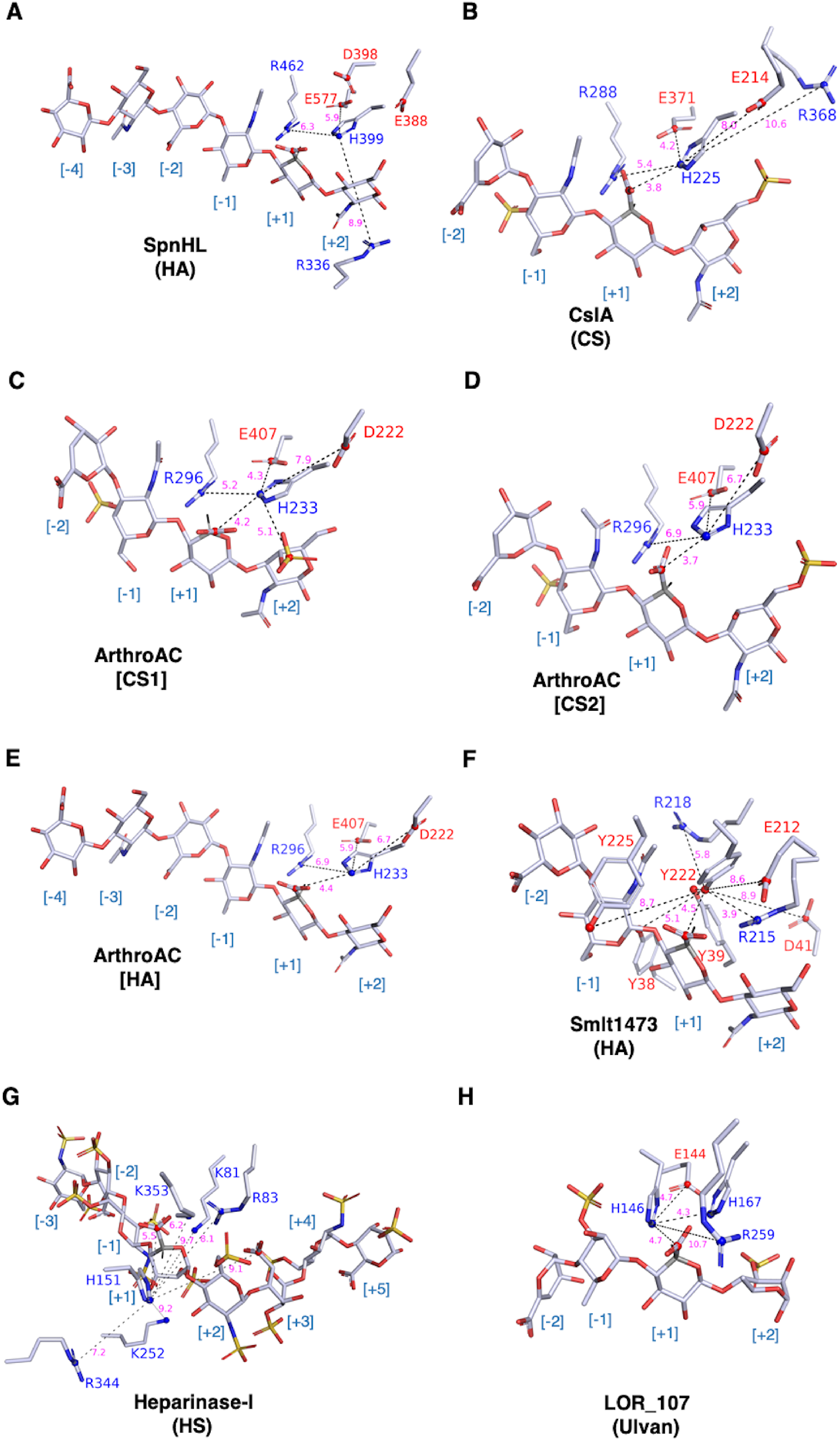
Electrostatic microenvironment around catalytic base of selected Poly-GlcA/IdoA lyases. (A) SpnHL complexed with hexa-HA [-4 to +2], catalytic base: H399 [+1], (B) CslA complexed with tetra-CS [-2 to +2], catalytic base: H225 [+1], (C) ArthroAC complexed with tetra-CS1 [-2 to +2], CS1 has sulphate group at C4 position of CS substrate at [+2] subsite, catalytic base: H233, (D) ArthroAC complexed (modelled) with tetra-CS2 [-2 to +2], CS2 has sulphate group at C5 position of CS substrate at [+2] subsite, catalytic base: H233, (E) ArthroAC complexed (modelled) with hexa-HA [-2 to +2], catalytic base: R171, (F) Smlt1473 complexed (modelled) with tetra-HA [-2 to +2], catalytic base: Y222, (G) Heparinase-I complexed with octa-HS [-3 to +5], catalytic base: H151, (H) LOR_107 complexed with tetra-Ulvan [-2 to +2], catalytic base: H146. The figure colouring and styling are same as FIGURE 5 (see figure legends).

In contrast to SpnHL, the CslA in apo state has only one pKa lowering determinant R288 at [+1] subsite for catalytic base H225. The pKa raising determinants include H-bond and coulombic interactions from E371 at [+1] subsite, and other coulombic interaction from E214 at [+2] subsite **(Table S4a)**. In the bound state, the catalytic base pKa lowering determinants apart from R288 include R368 at [+2] subsite. The pKa raising determinants include H-bond interaction with sugar ring oxygen atom and E371 at [+1] subsite, coulombic interaction from E371 via the carboxylic group at [+1] subsite, and E214 at [+2] subsite. It is evident from analysis of pKa that CslA catalytic base H225 experiences more negative charge interaction for both apo and bound state in comparison to SpnHL catalytic base H399. Thus, CS specific CslA has higher optimum pH than HA specific SpnHL **(Table S4a, FIGURE 6B)**.

Like CslA, the ArthroAC’s catalytic base H233 in apo state, faces only one pka lowering determinant, R296 at [-1, +1] subsite. The pKa raising determinant include main chain H-bond contribution from H233 itself, and coulombic interaction from D222, and E407 at [+1] subsite **(Table S4a)**. In CS bound state, the pKa lowering determinants are same as the apo state, R296. Similarly, the pKa raising determinants from apo state are maintained, D222, and E407. However, the position of the catalytic base in between carboxylic groups at [+1] subsite, and sulphate group at [+2] subsite, lead to pKa raising interactions which include H-bonds (i) from sugar ring oxygen atom and carboxylic group at [+1] subsite, and (ii) two H-bond interactions from the sulphate group at the [+2] subsite **(Table S4a, FIGURE 6C)**. This particular CS substrate raises the pKa of ArthroAC in the alkaline pH range which is much higher than its acidic optimum pH reported in literature. Thus, we modelled a CS substrate (CsIA) with varying position of sulphate group, and relaxed the modelled structure to re-calculate the pKa. With new calculations, the pKa lowering determinant is same as before, R296; apart from the usual pKa raising coulombic interaction from D222, and E407, the other pKa raising determinants include two H-bond interactions from carboxylic group oxygen at [+1] subsite **(Table S4a, FIGURE 6D)**. The pKa raising contribution of sulphate group at [+2] subsite is not possible due to its differed position on the sugar ring as compared to the original bound substrate. Thus, the catalytic base pKa of this structural model is consistent with observed acidic range of optimum pH. Moreover, the orientation of carboxylic groups is also not in an ideal position for titration to form coulombic interactions in the bound state. This result shows a slight variation in substrate structure can have drastic effect on determining the resulting catalytic pKa base. Finally, we modelled an HA substrate from SpnHl into ArthroAC coordinates, relaxed the model and subjected it to pKa calculation. In the HA bound state, the pKa lowering determinant at [+1] subsite is the same as others, R296. The pKa raising determinant include, D222, E407, and carboxylic group at [+1] subsite. The resulting pKa is consistent with the observed acidic optimum pH of activity **(Table S4a, FIGURE 6E)**.

### PLs binding GlcA at [+1] subsite, displaying (α/α)_n_ barrel fold

Smlt1473 is the first biochemically and structurally characterized pH-directed multifunctional PL, which displays (α/α)_n_ barrel fold, and optimally cleaves HA at pH 5.0. This pH optimum is similar to its PL-8 counterparts, which also utilize a His/Tyr catalytic system. However, the His/Tyr system in Smlt1473 lie in *anti*-position in comparison to *syn*-position of PL-8. Thus, there is possibility of more than one catalytic base candidate based on catalytic geometry. The catalytic base candidates include, R215 interacting with carboxylic group at [+1] subsite, H168 which also interact with carboxylic group and other polar atom in sugar ring at [+1] subsite, and Y222 which is in position to act as both catalytic base and acid. We subjected the HA docked Smlt1473 structure to pKa calculation to analyse the above possibilities.

In the apo state, H168 pKa is well below 5, due to its lack of solvent accessibility and coulombic interaction with only pKa lowering basic amino acid R215 at [+1] subsite **(Table S4b)**. In contrast, R215 has a pKa value greater than its intrinsic pKa value due to pKa raising H-bond interaction from Y222 at [+1] subsite, and coulombic interaction from Y39, Y115, Y222 at [+1] subsite, and Y38, Y225 at [-1] subsite. Additionally, the coulombic interaction from the acidic amino acids such as E212 at [+1] subsite, and D41 at [+2] subsite, further raises the pKa **(Table S4b)**. For Y222, the pKa raising coulombic interaction from D41, Y38, Y39, Y115, Y225, and D122 at [-1] subsite overwhelms the pKa lowering H-bond interaction from R215, and coulombic interaction from R215 and R218 **(Table S4b)**. In the bound state, the increase in H168 pKa is consistent with the acidic optimum pH of activity. The pKa increasing interaction includes H-bond interaction with glycosidic oxygen atom between [+2] and [+1] sugar unit and sugar ring oxygen atom at [+1] subsite. The coulombic interaction from the substrate carboxylic group at [+1] subsite and D111 at [+2] subsite further contribute to the rise in H168 pKa **(Table S4b)**. For R215, there is no pKa lowering interactions, so its pKa is much higher and therefore unable to acts as catalytic base **(Table S4b)**. Similarly, Y222 is also substantially altered with pKa raising determinants, and thus also precludes it acting as base at acidic pH **(Table S4b, FIGURE 6F)**. The amino acid residue H168 has a pKa consistent to act as base, however its *anti*-position with respect to labile C5-H at [+1] subsite precludes such a possibility. The other residues which are in *syn*-position to labile C5-H have very high pKa values. Thus, a more comprehensive analyses based on QM/MM methods may be required to fully elucidate this mechanism.

### PLs binding IdoA at [+1] subsite displaying β-jellyroll fold

Heparinase-I is the only biochemically and structurally characterized PL which cleaves the most negatively charged glycan (heparin sulphate). It utilizes a His/Tyr system as discussed above for poly-GlcA cleaving PLs. In particular, the Heparinase-I His/Tyr system is more similar to Smlt1473; however, in contrast to Smlt1473, the IdoA substrate’s labile C5-H at [+1] subsite is in *syn*-position with respect to H151, clearly establishing H151 as the catalytic base. In the apo state, H151 has a very low pKa due to no solvent accessibility, and availability of only pKa lowering determinants which include, R83, K353 at [+2, +1] subsite, and R344 at [-1] subsite **(Table S5)**. In the bound state, H151 pKa increases in comparison to the apo state due to interaction mainly from substrate which include H-bond interactions from glycosidic oxygen atom between [+2] and [+1] subsite, ring oxygen atom at [+1] subsite, and main chain oxygen atom of G152 at [+1] subsite. The pKa raising coulombic interactions include contribution from carboxylic groups at [+3] and [+1] subsite. Whereas the pKa lowering coulombic interactions include, K81, K252 at [+1] subsite, R344 at [+1, -1] subsite, R83, K353, and interaction with substrate N-group bonded with SO_4_ group at [+2] and [-1] subsite. The binding of highly negatively charged substrate does not raise the pKa significantly due to predominant pKa lowering interaction and no solvent accessibility **(Table S5, FIGURE 6G)**.

### PLs binding GlcA at [+1] subsite displaying β-propeller fold

Ulvan is a sulphated, non-glycan anionic polysaccharide that contain GlcA group bound by PLs at [+1] subsite. LOR_107 is a biochemically and structurally characterized ulvan lyase that cleaves glycosidic bond between GlcA and RG[3S]. H146 is thought to act as both the base and acid due to its spatial position with respect to bound substrate at [+1] subsite, whereas R259 acts as charge neutralizer. This is the first instance of among PLs where His is acting as base as well as acid **[27]**. In the apo state, H146 forms two pKa lowering coulombic interactions with H167, and R259 at [+1] subsite. The pKa raising interactions include H-bond and coulombic interaction from E144 at [+1] subsite. Despite higher contribution from the negatively charged interactions, the low solvent accessibility adds the pka lowering determinants to decrease the pKa one unit below their intrinsic value **(Table S5)**. In the bound state, the pKa lowering determinants for H146 are preserved as apo states, H167, and R259. Apart from the pKa raising H-bond, and coulombic interaction from E144, the other pKa raising contribution from substrate include H-bond interactions from cleavable glycosidic bond oxygen atom at [-1, +1] subsite, ring oxygen atom at [+1] subsite, and the coulombic interaction from carboxylic group at [+1] subsite. Similar to apo state, the lack of solvent accessibility of the catalytic base overwhelms the pKa raising negatively charged interaction to maintain similar pKa as apo state **(Table S5, FIGURE 6H)**.

### PLs binding ManA at [+1] subsite displaying (εξ/εξ)_n_ barrel fold

The PL-5 family of PLs are biochemically and structurally well characterized for cleaving ManA at optimum alkaline pH values. The selected PL-5 enzymes for pKa analysis include alginate lyase A1-III and Smlt1473; the catalytic system utilized by these PLs is similar to heparinase-I as discussed above. However, the alginate lyase A1-III, and Smlt1473 have structurally more than one choice for catalytic base residue due to their catalytic geometry. These residues include, H192, R239, and Y246 for alginate lyase A1-III, and structurally conserved H168, R215, and Y222 for Smlt1473. We analysed each of these amino acid residues for their ability to act as the catalytic base.

For alginate lyase A1-III, the H192 in apo state interacts with pKa lowering R196 and R239 occupying [+2], and [+1] subsite respectively. The pKa raising determinants include H-bond interaction with E140, and coulombic interaction with E140, and E236 at [+1] subsite. A very low H192 solvent accessibility balance the predominant negatively charged interaction resulting in pKa rise slightly above its intrinsic value **(Table S6)**. In the bound state, H192 maintains same pKa lowering residues as apo state, R196, and R239. The pKa raising interactions are also maintained as apo state, E140, and E236. However, the additional pKa raising contribution from bound substrate (sugar ring oxygen, and carboxylic group) at [+1] subsite negate the pKa lowering effect of H192’s lack of solvent accessibility. Interestingly, the pKa increases to values consistent with the optimum pH of activity **(Table S6)**. For R239 in apo state, there are only weak but pKa raising determinants at [+1] subsite which include H-bond interaction with Y246, and coulombic interaction involving Y137, Y246, E236, and E241 at [+1] subsite. The high solvent accessibility of R239 balance the contribution of pKa lowering interactions resulting in intrinsic pKa values **(Table S6)**. In the bound state, R239 interacts weakly with pKa lowering R67 at [-1] subsite. The pKa raising interactions from apo state are preserved except for new contributions from Y68 and substrate carboxylic group at [+1] subsite. The pKa raising interactions in the bound state overwhelm the pKa lowering effect of R239 low solvent accessibility, precluding its possibility as catalytic base, and thus limit its use as an aide in only charge neutralization and substrate stabilization **(Table S6)**. For Y246 in apo state, the pKa lowering determinants include H-bond and coulombic interaction from R239 at [+1], and coulombic interaction from R306 at [-2] subsite. The pKa raising contribution in form of coulombic interactions include E236, E241 at [+1] subsite, and D304, and D314 at [-2] subsite **(Table S6)**. For Y246 in bound state, the pKa lowering interactions from R239 and R306 are preserved, but include additional contribution from H-bond interactions from substrate glycosidic oxygen at [+1, -1] subsite, and hydroxyl group oxygen R67 at [-1] subsite. The pKa raising contributions from negatively charged residues in apo state are preserved except D314. The new contributions include H-bond and coulombic interactions from Y68, and coulombic interaction from Y80 at [-1] subsite with a strong, moderate, and almost negligible coulombic interaction from substrate carboxylic groups at [+1], [-1], and [-2] subsites, respectively. The lack of solvent accessibility of Y246 heavily aids the pKa raising contribution from negatively charged residues, and thus pKa jumps well above the intrinsic values **(Table S6, FIGURE 7A)**. This present a situation similar to what we observed above for pKa analysis of Smlt1473 docked with HA, the only residue consistently acting as catalytic base (H168) at the observed optimum pH of activity, which structurally is similar to the conserved His (H192) residue. However, its *anti*-orientation **[1, 7-9]** with substrate’s labile C5-H group at [+1] subsite geometrically preclude its ability to act as catalytic base.

**FIGURE 7.**
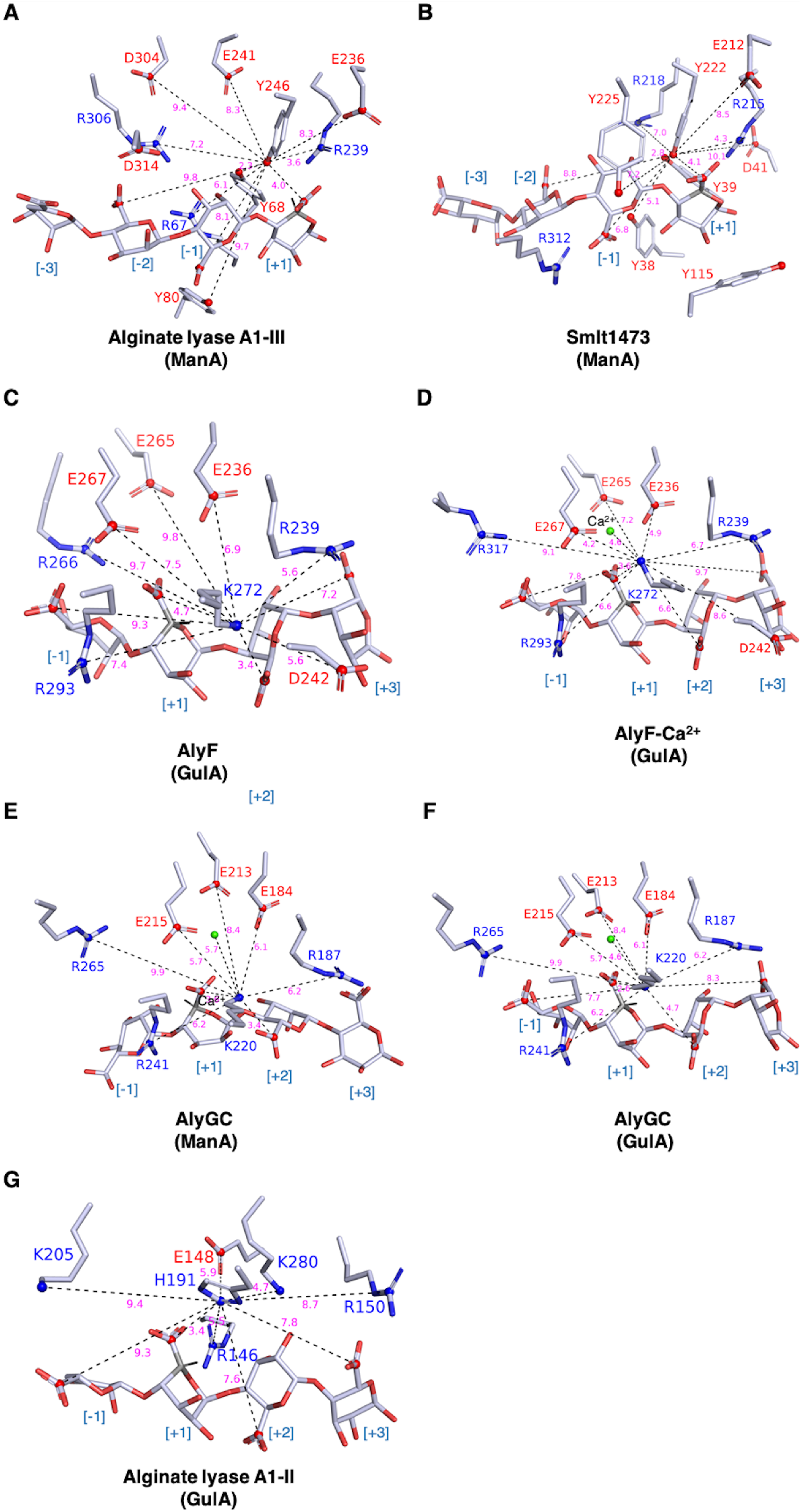
Electrostatic microenvironment around catalytic base of selected Poly-ManA/GulA lyases. (A) Alginate lyase A1-III complexed with tetra-ManA [-3 to +1], catalytic base: Y246 [+1], (B) Smlt1473 complexed with tetra-ManA [-3 to +1], catalytic base: Y222 [+1], (C) AlyF complexed with tetra-GulA [-1 to +3], catalytic base: K272 [+1], (D) AlyF complexed with Ca^2+^ (modelled), and tetra-GulA [-1 to +3], catalytic base: K272 [+1], (E) AlyGC complexed with Ca^2+^, and tetra-ManA [-1 to +3], catalytic base: K220, (F) AlyGC complexed (modelled) with Ca^2+^, and tetra-GulA [-1 to +3], catalytic base: K220, (G) Alginate lyase A1-II complexed with tetra-GulA [-1 to +3], catalytic base: H191 [+1]. The figure colouring and styling are same as FIGURE 5 (see figure legends).

For pKa analysis of ManA-bound Smlt1473, we modelled the sugar unit at [-3] subsite as it is not modelled in the deposited crystal structure. In the bound state, H168 has one pKa lowering determinant, R215 at [+1] subsite. In contrast, the pKa raising determinants include H-bond interaction with sugar ring oxygen at [+1] subsite. The other pKa raising determinant includes D111 and carboxylic group at [+2], and [+1] subsites, respectively. The lack of solvent accessibility for H168 lowers the pKa well below the intrinsic values **(Table S6)**. For R215 in the bound state, there is the presence of only pKa raising interactions which include H-bond and coulombic interaction with carboxylic group at [+1] subsite, and Y222 at [+1, -1] subsite. The other pKa raising determinants include D41(93) at [+2] subsite, Y39(27), Y115(17), E212(0) at [+1] subsite, and Y38, Y225 at [-1] subsite. The presence of only negatively charged interactions balance the pKa lowering effect for R215 due to low solvent accessibility, thus rendering it to function only as a substrate stabilizer **(Table S6)**. For Y222, the pKa lowering determinants include H-bond interaction with glycosidic oxygen at [+1, -1] subsite, C2-hydroxyl group at [-1] subsite, and R215 at [+1] subsite. The pKa lowering coulombic interactions are from side chains of R215 and R218. The pKa raising determinants include H-bond interaction from Y39 and coulombic interaction with D41, E212, Y38, Y39, Y225 and carboxylic groups at [+1, -1, -2] subsites **(TABLE S6, FIGURE 7B)**.

The lack of solvent accessibility coupled with overwhelming interactions from negatively charged determinants from protein and substrates significantly raise Y222 pKa to act as the catalytic base **(Table S6)**. Interestingly, only H168 has a suitable electrostatic environment for functioning as catalytic base, but its *anti-*orientation with respect to labile substrate C5-H at [+1] subsite geometrically precludes that from happening. In previous studies, It has been confirmed that Smlt1473 can cleave poly-GulA (C5 epimer of ManA) **[2, 3]**, which contains labile substrate C5-H in *syn*-orientation with respect to H168. However, Smlt1473 activity toward poly-GulA is low, suggesting an elusive catalytic mechanism not yet captured by current structural and computational analysis. In subsequent sections, we end this structural-computational analyses with discussion of PLs binding GulA at [+1] subsite, where we will encounter PL-6 with parallel β-helix fold which are structurally and catalytically similar to poly-GalA lyase discussed in previous section. We will also encounter PL-7 family of poly-GulA lyase utilizing a His/Tyr system similar to PL-5, but embedded in a β-sandwich fold.

### PLs binding GulA at [+1] subsite displaying parallel β-helix fold

The PL-6 family of PLs are known to cleave alginates and chondroitin sulphates, and like their structural homolog poly-GalA lyase, they utilize metal ion (Ca^2+^) for charge neutralization. The only biochemically and structurally well-characterized (apo and substrate bound) PL-6 are poly-GulA specific lyase, AlyF, and AlyGC; hence we selected these two PLs for structural and computational analyses. It has been reported for AlyF that, it does not require the presence of Ca^2+^ ion for its enzyme catalysis **[29]**. However, it does contain the Ca^2+^ ion binding site like other PL6 (e.g. AlyGC), so we added the Ca^2+^ ion at the identified binding site in the substrate bound structure, and relaxed it before pKa analysis. First, we will discuss the result of AlyF’s pKa analysis without bound Ca^2+^ as reported in literature **[29]**. In apo state, K272’s pka lowering determinants include, R239 at [+3] subsite, R266, R293 at [-1] subsite. The pKa raising determinants include, D242 at [+3] subsite, and E265, E236, E267 at [+1] subsite. The higher contribution of negatively charged pKa raising determinants is balanced largely by the lack of solvent accessibility of K272, thus pKa decreases by only by a unit below the intrinsic value. The protein coordinate in apo state does not contain the Ca^2+^ ion at [+1] subsite, or we otherwise would have capture lower pKa values **(Table S7)**. In bound state, the pKa lowering determinants are preserved as apo state, and have the same low solvent accessibility. The pKa raising determinants apart from D242, E265, E236, and E267 include H-bond and coulombic interaction from carboxylic group at [+2] subsite, and other coulombic interaction from carboxylic groups at [+3], [+1], and [-1] subsite. Interestingly, the pKa values of all pKa raising determinants, except the substrate carboxylic group and D242, is much higher than their intrinsic values, and lie in near neutral to alkaline pH range (E265, E236, and E267). Neglecting the contribution of these determinants lowers the K272 pKa value in the bound state to values similar to apo AlyF **(Table S7, FIGURE 7C)**. We will next discuss the result of pKa analysis of substrate bound AlyF in the presence of Ca^2+^ ion modelled at the [+1] subsite, as this may shed light on why AlyF may not require Ca^2+^ for its functioning. K272 pKa lowering determinants apart from R239 and R293 include R317 at [-1] subsite. R317 replaces nearby R266 from previously discussed substrate bound state in AlyF without Ca^2+^ ion. The pKa raising determinants are same as discussed previously, but in contrast to AlyF (without Ca^2+^) the pKa of residue E265 is in the alkaline range. The lowering of pKa for other residues such as E236 and E267 can be attributed to the bound Ca^2+^ ion, as its presence has repacked the side chain of E236 and E267 in such a way that they experience less pKa raising negative charge interactions in comparison to their counterpart in substrate bound AlyF without Ca^2+^ ion. This suggest that the presence of Ca^2+^ ion lowers the pKa of acidic amino acid cluster at [+1] subsite. The low pKa of these amino acids will make them negatively charged at very low pH, and resulting negative charge interaction will raise the catalytic base pKa. The Ca^2+^ ion should lower the catalytic base pKa, but overwhelming negative charge from substrate carboxylic group and acidic amino acid cluster at [+1] subsite prevent the decrease in pKa **(Table S7, FIGURE 7D)**.

Next, we performed the pKa calculation for AlyGC which is known to utilize Ca^2+^ ion for catalysis, and we have compared it with the AlyF’s active site pKa determinants to understand the differences in the catalytic mechanism. For AlyGC’s catalytic base K220 in apo state, the pKa lowering determinants apart from its low solvent accessibility include positively charged coulombic interaction from bound Ca^2+^ ion at [+1] subsite, R187 at [+3], and R241 at [+1, -1] subsite. The pKa raising determinants include negatively charged coulombic interactions from E184 at [+2] subsite, E213, and E215 at [+1] subsite. The number of both pKa lowering as well as raising residue interaction with catalytic base is less compare to AlyF apo structure **(Table S7)**. The GulA bound state for AlyGC was obtained by modelling the GulA coordinate obtained from atomic superposition of AlyGC (complexed with ManA-Ca^2+^) with AlyF and structure relaxation using Rosetta. The pKa lowering interaction for K220 apart from Ca^2+^, R187, and R241 include, weakly interacting R265 at [-1] subsite. In contrast, the pKa raising determinants apart from E184, E213, and E215 include H-bond and coulombic interaction from carboxylic group at [+1] subsite. The other coulombic interactions include carboxylic group from [+3, +2] subsites. The carboxylic group interaction is also less in comparison to bound AlyF structure **(Table S7, FIGURE 7F)**. The ManA bound crystal structure also have same number of pka lowering and raising determinants except for the carboxylic group. The change in carboxylic group contribution can be attributed to change in ManA’s substrate pose in contrast to its C5-epimer GulA **(Table S7, FIGURE 7E))**. Recently, the PLs from this family (PL-6) has also been implicated in cleavage of ManA containing substrate **[30]**. However, the catalytic Lys is in opposite direction of ManA’s C-5 orientation at [+1] subsite suggesting a highly dynamic binding site like their PL-5 counterpart.

### PLs binding GulA at [+1] subsite displaying β-sandwich fold

The PL-7 family of PLs are known to specifically bind and cleave poly-GulA. Unlike PL-6’s Lys/Arg catalytic system, the alginate lyase from PL-7 utilize His/Tyr system similar to PL-5 PLs. Alginate lyase A1-II is one such structurally and biochemically characterized PL which we have selected for pka analyses. The catalytic base H191, interacts predominantly with pka lowering basic amino acids which include, R150 at [+3] subsite, K280 at [+2] subsite, R146 at [+1] subsite, and K205 at [-1] subsite. The pka raising determinants include, backbone H-bond interaction from G192 at [+1] subsite, and coulombic interaction from E148 at [+1] subsite **(Table S7)**. In bound state the pka raising determinants are same as apo state, R146, R150, K280, and K205. The pka raising contribution include coulombic interaction contribution from E148, and carboxylic group at [+3, +2, +1, -1] subsite. The increase in negative charge contribution from substrate carboxylic group raise the pKa of H191 in comparison to apo state **(Table S7, FIGURE 7G)**.

### Conclusions

The Smlt1473’s unique ability to cleave multiple substrates as a function of pH inspired the preliminary analyses and compiling of data in light of the PL-pH functional relationship (Table 1). The finding that mono-specific PLs utilize similar pH as Smlt1473 does for each of its multiple substrates extended our search for all major substrates and focused explicitly on substrate-pH pairing (FIGURE 2-4). Interestingly, huge structural differences among PLs did not deter catalytic pH range for the same substrates. The PLs among each substrate group (FIGURE 1, S1) and members within and among families were structurally and sequentially compared to account for similarities and differences. For PLs with closely related structures, both sequence and structural alignments were performed. We also found that substrate-coordinate-based structural alignment is more informative. They can be used to find structural convergence among PLs, which fundamentally vary at the fold level (FIGURE S4-13). The result of PLs subjected to pKa analysis was not only based on numerical values which set very close arguments with the observed working pH of PLs but were also based on the coulombic and other non-covalent interaction data (which influence numerical values) modelled in stick representation (FIGURE 5-7). Overall, this study aims to provide a knowledge base to inspire the development of future bio-engineering research involving PLs, or other enzymes where pH plays a significant role in substrate binding or cleavage. However, further analyses of change in charged surfaces area [electropositive (low pH) 🡺 electronegative (high pH)] among PLs by modelling protein-wide coulombic interaction network; and, comparison with an electrostatic map at different pH (including optimum pH as centre) will generate further information which can be used to identify region distal to the active site for controlling modelling new native or non-native functions.

## Materials and methods

### Selection of PLs for the structural and computational analyses

The PLs tabulated in Table S1 are biochemically, and structurally characterized representatives of their family. For this study, we have selected PLs which are both enzymatically and structurally characterized in apo as well as substrate-bound form. We have to make sure that, the PLs selected for each substrate group cover all possible optimum pH ranges within the substrate-specific pH range of activity. For proper structural, and computational analyses, the mutated PL-substrate complex is modelled *in silico* into wildtype by Rosetta relax protocol. Bad structures, for e.g. PDB file containing multiple charges changing mutations, protein sequence from PDB varying significantly from Uniprot sequence, improperly build ligands, and protein structures have not been selected. The PL selected by the above criteria is tabulated (see Table 1).

### Rosetta’s Fast relax protocol

Use of active site mutations (Arg/His/Lys to Ala/Leu; Tyr to Phe) is generally used to complex the substrate in the crystal structure. For structural analysis, and computational pKa calculation, we have dis-mutated the crystal structures *in silico* which may result in a vdW clash with the substrate. To avoid such a clash, we use Rosetta’s fast relax protocol [31]. The usage information, script and protocol can be found at Rosetta Commons (https://www.rosettacommons.org).

### pk_a_ analyses

For both ‘apo’ and ligand-bound structure, we have performed pk_a_ calculation at optimum pH and used the command-line version of PROPKA31 which incorporate ligands and ions into the calculations [32, 33]. The result of the pk_a_ calculation has been used to (1) generate H-bond, coulombic interaction network for the catalytic base, and (2) to model negative and positive charged amino acid clusters around the active site.

## Supplementary material description

The supplementary material can be found in the pdf file Supplementary_Fig_and_Tables_SP_BWB_RA_14Feb23.

## Author Contributions

S.P., Conceptualization, Methodology, Data curation, Software, Formal analysis, Validation, Writing - original draft, Writing - review & editing.

B.W.B., Funding acquisition, Investigation, Writing - review & editing, Supervision R.A., Conceptualization, Methodology, Formal analysis, Supervision, Funding acquisition, Investigation, Project administration, Writing - original draft, Writing - review & editing, Resources

## Author Approvals

All authors have seen and approved the manuscript, and that it hasn’t been accepted or published elsewhere

## Acknowledgments

S. P., and R. A: We thank NISER for funding. This work supported in part by Department of Biotechnology, Govt. of India grant number BT/PR15324/BRB/10/1482/2016.

B. W. B: This work was supported in part by a grant from the Center for Innovative Technology, a Research Innovation Award from the University of Virginia School of Engineering and Applied Science and a grant from the National Science Foundation (CBET 1452855).

