## Supplementary for "Structural analyses of ‘substrate-pH of activity’ pairing observed in Polysaccharide lyases"

**Note S1: The general $pk_a$ perturbing factors for titrable side chains in protein molecule.**

The side chain of titrable amino acid can attain different charge state under varying pH values; and such amino acids include- (i) acidic amino acids (negatively charged, Asp, Glu, Cys, and Tyr), and (ii) basic amino acids (positively charged, His, Lys, and Arg) (ref). The pH at which the titrable amino acid side chain attain charge and uncharged state equilibrium is called $pk_a$. In fully solvated state, the $pk_a$ values are known as intrinsic values; and are calculated by the help of NMR spectroscopy of model peptides containing the titrable amino acid of choice (ref). The typical intrinsic $pk_a$ values of Asp, Cys, Glu, His, Lys, Arg, and Tyr are 3.9, 8.3, 4.3, 6.0, 10.8, 12.5, and 10.9 respectively (ref). However, the proteins are dynamic molecule with chemically varied amino acid side chains in close proximity and variable solvent accessibility, the intrinsic $pk_a$ values does not hold ground. Thus, the $pk_a$ values varies depending upon the type of amino acid side chain, percent solvent accessibility surface area (Born effect), and inter or intramolecular interactions (Charge-charge, and charge-dipole interactions) (ref). However, unlike small peptide and proteins, the large and more chemically complex proteins are not amenable to $pk_a$ calculation by NMR spectroscopy. Thus, we have utilized the computational $pk_a$ calculation (see, method section of main article) which weigh in the above mentioned $pk_a$ perturbing (or varying) factors. A brief postulates of these $pk_a$ perturbing factors and their effect on various group of amino acids are as under-

- **Born effect:** When titrable amino acids are partially or completely buried the neutral state of their ionizable groups are favoured as it is hard to transfer charges in low di-electric environment of protein interior. Consequently, pKa of Asp, Glu, Cys, and Tyr is raised, and that of His, Lys, and Arg is lowered.
- **Charge-charge interactions:** The pKa of all titrable amino acid will be lowered by the positively charged environment, and raised by the negatively charged environment. At pH=pKa, the ionizable group of Asp, Glu, Cys, and Tyr will have net charge of -1/2, and that of His, Lys, and Arg will have net charge of +1/2. For Asp, Glu, Cys, and Tyr, the condition pH<pKa and pH>pKa will respectively lead to neutrality and negative charge of ionizable groups. Opposite effect is observed for His, Lys, Arg where the condition pH<pKa and pH>pKa lead respectively to positive charge and neutrality of ionizable group.
- **Charge-dipole interactions:** If such interactions are more favourable with protonated state of ionizable group the pKa will be raised, whereas similar interaction with deprotonated state of ionizable group will lowers the pKa. This imply that, introduction of such interactions to Asp, Glu, Cys, and Tyr in their protonated state (pH<pKa) will raise their pKa values and vice versa. On the other hand, such interaction formed by His, Lys, and Arg in their deprotonated form (pH>pKa) will lowered their $pK_a$ and vice versa.

Supplementary Information

**Table S1: Census of structurally and functionally characterized PLs from CAZY database**

| Substrate type | Substrate | Catalyzing PL-family | Subfamily | E.C. No. | Organism | PDB id (apo / substrate bound) | Opt pH of activity |
|---|---|---|---|---|---|---|---|
| **Galacturonic acid (GalA)** | Pectate | PL-1 | 5 | 4.2.2.2 | *Bacillus* sp. NI6-5 | 3VMV (pH-8.0)/3VMW (tri-GalA, pH-8.5) | 11.5 |
| | Pectate | PL-1 | 6 | 4.2.2.2 | *Bacillus subtilis* subsp. *Subtilis* str. 168 | 1BN8 (pH-6.5)/2O17(hexa-sach., pH-4.6) | 8.5 |
| | Pectate | PL-1 | 3 | 4.2.2.2 | *Dickeya chrrysanthemi* EC16 | 1O8F (pH9.5)/2EWE (penta-GalA, pH-9.5) | 9.5 |
| | Pectate | PL-1 | - | 4.2.2.2 | *Thermotoga maritima* MSB8 | 3ZSC(tri-sach, pH-4.0) | 9 |
| | Pectate | PL-2 | 1 | 4.2.2.2 | *Yersinia enterocolitica* subsp. *enterocolitica* 8081 | 2V8J(Mn2+)/2V8K (tri-GalA) | 9.61 |
| | polyGalA | PL-2 | 2 | - | *Vibrio vulnificus* YJ016 | 5A29 (pH-7.0) | 9.3 |
| | Pectate | PL-3 | 1 | 4.2.2.2 | *Bacillus* sp. KSM-P15 | 1EE6 (pH-6.7) | 10.5 |
| | Pectate | PL-3 | 1 | 4.2.2.2 | *Caldicellulosiruptor bescii* DSM 6725 | 3TG9 (pH-8.5)/4Z03 (tri-GalA, pH-7.7) | 8.5 |
| | Pectate | PL-3 | 5 | 4.2.2.2 | *Dickeya dadantii* 3937 | 3B4N (pH-6.5)/3B8Y (pH-6.5) | 9.2 |
| | Rhamnogalctouronate | PL-9 | 1 | 4.2.2.23 | *Bacteroides thetaiotaomicron* VPI-5482 | 5OLQ/5OLOS (tri-GalA) | unconfirmed |
| | Pectate | PL-9 | 1 | 4.2.2.2 | *Dickeya dadantii* 3937 | 1RU4 (pH-5.6) | pH-8.0 |
| | Pectate | PL-10 | 1 | 4.2.2.2 | *Cellvibrio japonicus* Ueda107 | 1GXM (pH-5.2)/1GXO (tri-GalA, pH-5.2) | 10.3 |
| | Pectate | PL-10 | 1 | 4.2.2.2 | *Niveispirillum irakense* KBC1 | 1R76 (pH-7.8) | 9 |
| | Oligogalacturonate | PL-22 | 1 | 4.2.2.6 | *Yersinia enterocolitica* subsp. *enterocolitica* 8081 | 3PE7 | 7.3-7.7 |
| **Glucuronic acid (GlcA)** | Hyaluronate | PL-8 | 1 | 4.2.2.1 | *Streptococcus pnemoniae* R6 | 1EGU (pH-6.0)/1LOH (3HA, pH-6.0) | 6 |
| | Hyaluronate | PL-8 | 1 | 4.2.2.1 | *Steptococcus agalactiae* NEM316 | 1F1S (pH-6.0)/1LXM (3HA, pH-6.0) | 5 |
| | Hyaluronate | PL-8 | | 4.2.2.1 | *Streptomyces coelicolor* | 2WCO (1HA) | 5.2 |
| | Chondroitin sulfate | PL-8 | 3 | 4.2.2.5 | *Pedobacter heparinus* DSM 2366 | 1CB8 (pH-8.0)/1HMW (tetra-CS, pH-7.5) | 8 |
| | Chondroitin sulfate / Hyaluronate | PL-8 | | 4.2.2.1 / 4.2.2.5 | *Paenarthrobacter aurescens* | 1RWA (pH-6.4)/1RWF (iduronated tri-CS, pH-6.4) | 6.0 (more activity against HA) |
| | Chondroitin sulfate | PL-23 | | 4.2.2.- | *Autographa californica* nucleopolyhedrovirus | 3VSM (pH-8.0) | 4.0-9.0 |
| | Ulvan | PL-24 | | 4.2.2.- | *Alteromonas* sp. LOR | 6BYP (pH-6.5)/6BYT (tetra-sach, pH-6.5) | 7.5 (Type-A Ulvan lyase)* |
| **Iduronic acid (IdoA)** | Chondroitin B* | PL-6 | 1 | 4.2.2.19 | *Pedobacter heparinus* DSM 2366 | 1DBG (pH-8.6)/1OFM (tetra-CS, pH-8.7 ) | 8 |
| | Heparan sulfate, heparin | PL-12 | 2 | 4.2.2.7 / 4.2.2.8 | *Bacteroides thetaiotaomicron* VPI-5482 | 4FNV | 6.7-7.3 |
| | Heparan sulfate, heparin | PL-12 | 2 | 4.2.2.7 / 4.2.2.8 | *Bacteroides thetaiotaomicron* VPI-5482 | 5JMF (pH-5.5) | 7.5 |
| | Heparan sulfate | PL-12 | 2 | 4.2.2.8 | *Pedobacter heparinus* DSM 2366 | 4MMH (pH-7.0) | 7.5 |
| | Heparin | PL-13 | _ | 4.2.2.7 | *Bacteroides thetaiotaomicron* VPI-5482 | 3IKW (pH-7.5)/3INA (dodecasaccharide, pH-5.5) | 7.5 |
| | Heparan sulfate, heparin | PL-21 | 1 | 4.2.2.7 / 4.2.2.8 | *Pedobacter heparinus* DSM 2366 | 2FUQ (pH-5)/3E7J (tetra-heparan sulfate | 7 |
| **Mannuronic acid (ManA)** | Mannuronate | PL-5 | | 4.2.2.3 | *Sphingomonas* sp. A1 | 1QAZ/4F10, 4F13 (4ManA) | 8 |
| | Mannuronate | PL-5 | 1 | 4.2.2.3 | *Pseudomonas aeruginosa* PAO1 | 4OZW/4OZV | 8 |
| | Mannuronate | PL-5 | | 4.2.2.3 | *Stenotrophomonas maltophilia* K279a | to submit (manuscript_1) | 9 |
| | Mannuronate | PL-31 | | 4.2.2.3 | *Paenibacillus* sp. FPU-7 | 6KFN (pH-6.5) | 7.0-7.5 |
| | Mannuronate | PL-7 | | 4.2.2.3 | *Flavobacterium* sp. UMI-01 | 5Y33 (pH-7.0) | 6.8-8.0 |
| **Guluronic acid (GulA)** | Guluronate | PL-6 | 1 | 4.2.2.11 | *Paraglaciecola chathamensis* S18K6T | 5GKD (pH-8.5)/5GKQ ( tetra-ManA, pH-8.5) | 7 |
| | Guluronate | PL-6 | | 4.2.2.11 | *Vibrio splendidus* | 6ITG (pH-7.0)/6A40(tetra-GulA, pH-7.5) | 7.5 |
| | Guluronate | PL-7 | | 4.2.2.- | *Sphingomonas* sp. A1 | 2CWS/2ZAC (4Alg) | 7.5 |
| | Guluronate | PL-7 | 3 | 4.2.2.11 | *Zobellia galactanivorans* DsijT | 3ZPY (pH-7.0) | 7 |

Supplementary Information

#### Table S2

Table S2: The distribution of catalytic base's pka determinants at active site of GalA cleaving PLs belonging to family PL-1

| Protein name | PL family | Optimum pH | Catalytic base | pka raising determinants | pka lowering determinants | Subsites | pka change due to desolvation |
|---|---|---|---|---|---|---|---|
| **BsPel (apo)** | PL-1 (subfamily-6) | 8.5 | R279 (%SASA=6, pka = 11.95) | D173 (%SASA=64, 0.12)*, D246 (%SASA=3, 0.32), Y308 (%SASA=22, 0.51) | | [+2] | -2.21 |
| | | | | | Ca2+ (-1.01), R284 (%SASA=58, -0.14) | [+1] | |
| | | | | D184 (%SASA=0, 0.14), D223 (%SASA=0, 0.93), D227 (%SASA=0, 0.78) | | [-1] | |
| **BsPel (substrate bound)** | | | R279 (%SASA=0, pka = 7.60) | D173 (%SASA=43, 0.21), D246 (%SASA=0, 0.55), Y308 (%SASA=9, 1.05), substrate's COO- group (%SASA=54, 0.47) | R282 (%SASA=29, -0.41) | [+2] | -3.46 |
| | | | | | Ca2+ (-2.03), R284 (%SASA=41, -0.14) | [+1] | |
| | | | | D223 (%SASA=0, 0.61), D227 (%SASA=0, 0.42), substrate's COO- group (%SASA=14, 0.37) | Ca2+ (-2.02), Ca2+ (-0.53) | [-1] | |
| **PelC (apo)** | PL-1 (subfamily-3) | 9.5 | R218 (%SASA = 20, pKa = 11.75) | Y268 (%SASA=57, 0.04) | | [+3] | -1.68 |
| | | | | D162 (%SASA=55, 0.46) | | [+2] | |
| | | | | | Ca2+ (-0.36), R245 (%SASA=38, -0.14) | [+1] | |
| | | | | D131 (%SASA=3, 0.04), E166 (%SASA=17, 0.33), D170 (%SASA=0, 0.56) | | [-1] | |
| **PelC (substrate bound)** | | | R218 (%SASA = 0, pKa = 7.24) | Y268 (%SASA=35, 0.07), substrate's COO- group (%SASA=30, 0.68) | | [+3] | -3.71 |
| | | | | D162 (%SASA=53, 0.37), E154 (%SASA=51, 0.02), D160 (%SASA=95, 0.05), substrate's COO- group (%SASA=28, 0.78) | Ca2+ (-1.74) | [+2] | |
| | | | | substrate's COO- group (%SASA=38, 1.90) | Ca2+ (-3.48), R223 (%SASA=28, -0.25), R245 (%SASA=14, -0.25) | [+1] | |
| | | | | D131 (%SASA=0, 0.04), E166 (%SASA=3, 0.65), D170 (%SASA=0, 0.63), substrate's COO- group (%SASA=31, 0.27) | Ca2+ (-0.51), Ca2+ (-0.77) | [-1] | |
| **PelA (apo)** | PL-1 (subfamily-5) | 11.5 | R207 (%SASA=16, pka = 11.79) | Y269 (%SASA=22, 0.05) | | [+3] | -1.98 |
| | | | | | | [+2] | |
| | | | | D150 (%SASA=37, 0.12) | R212 (%SASA=46, -0.19) | [+1] | |
| | | | | D153 (%SASA=37, 0.41), D157 (%SASA=0, 0.88) | | [-1] | |
| **PelA (substrate bound)** | | | R207 (%SASA=7, pka = 11.24) | Y269 (%SASA=12, 0.20) | | [+3] | -3.47 |
| | | | | substrate's COO- group (%SASA=39, 0.59) | | [+2] | |
| | | | | D150 (%SASA=82, 0.17), substrate's COO- group (%SASA=53, 1.6) | R212 (%SASA=46, -0.19) | [+1] | |
| | | | | D153 (%SASA=18, 0.41), D157 (%SASA=0, 0.88), substrate's COO- group (%SASA=38, 0.18) | Ca2+ (-0.32), Ca2+ (-0.99) | [-1] | |

#### Table S3

Table S3: The distribution of catalytic base's pka determinants at active site of GalA cleaving PLs belonging to family PL-2, 3, and 10

| Protein name | PL family | Optimum pH | Catalytic Base | pka raising determinants | pka lowering determinants | Subsites | pka change due to desolvation of catalytic base |
|---|---|---|---|---|---|---|---|
| **PecB (apo)** | PL-3 | 8.5 | K108 (%SASA=51, pka=9.61) | | K130 (%SASA=58, -0.17), K160 (%SASA=93, -0.01) | [+3] | -1.17 |
| | | | | D107(%SASA=63, 0.39) | R133 (%SASA=59, -0.22) | [+2] | |
| | | | | | Ca2+ (-0.36) | [+1] | |
| | | | | E39($\%_{SASA}$=100, 0.03), D64($\%_{SASA}$=31, 0.05), E84($\%_{SASA}$=68, 0.09), D85($\%_{SASA}$=25, 0.49) | | [-1] | |
| **PecB (substrate bound)** | | | K108 (%SASA=51, pka=7.66) | C157($\%_{SASA}$=30, 0.11), D162 ($\%_{SASA}$=78, 0.04), and E183($\%_{SASA}$=87, 0.0) | K130 (%SASA=43, -0.81), K160 (%SASA=75, -0.06) | [+3] | -3.26 |
| | | | | D107(%SASA=45, 0.99), substrate's COO- group (%SASA=81, 1.08) | R133 (%SASA=43, -0.45) | [+2] | |
| | | | | substrate's COO- group (%SASA=68, 0.52) | Ca2+ (-0.80) | [+1] | |
| | | | | E39($\%_{SASA}$=75, 0.10), E84($\%_{SASA}$=41, 0.19), D85($\%_{SASA}$=0, 0.10), substrate's COO- group (%SASA=47, 0.13) | Ca2+ (-0.60), Ca2+ (-0.11) | [-1] | |
| **PL2A (apo)** | PL-2 | 9.6 | R171 (%SASA=45, pka = 11.30) | | | [+2] | -1.63 |
| | | | | E130($\%_{SASA}$=25, 0.28) | | [+1] | |
| | | | | E515($\%_{SASA}$=66, 0.02), Y235($\%_{SASA}$=0, 0.02), Y248($\%_{SASA}$=37, 0.32), Y364($\%_{SASA}$=63, 0.06) | $Mn^{2+}$ (-0.28) | [-1] | |
| | | | | E302($\%_{SASA}$=25, 0.01) | | [-2] | |
| **PL2A (substrate bound)** | | | R171 (%SASA=4, pka=11.43) | E126($\%_{SASA}$=54), Y162($\%_{SASA}$=31) | | [+2] | -3.41 |
| | | | | E130($\%_{SASA}$=25, 0.28), E287(%SASA=0), substrate's COO- group (%SASA=0, 2.22) | $Mn^{2+}$ (-2.58), R272(%SASA=8, -0.30) | [+1] | |
| | | | | E515($\%_{SASA}$=66, 0.02), Y248($\%_{SASA}$=37, 0.32), Y364($\%_{SASA}$=63, 0.06), substrate's COO- group (%SASA=16, 0.56) | | [-1] | |
| | | | | | | [-2] | |
| **Pel10A (apo)** | PL-10 | 10.3 | R524 (%SASA=53, pka = 14.65) | Y443($\%_{SASA}$=61, 0.41) | | [+2] | -1.14 |
| | | | | E527($\%_{SASA}$=22, 2.34), D389(%SASA=38, 0.23), Y526(%SASA=75, 0.54) | | [+1] | |
| | | | | Y629($\%_{SASA}$=10, 0.52), D451(%SASA=35, 0.04) | $Ca^{2+}$ (-0.80) | [-1] | |
| **Pel10A (substrate bound)** | | | R524 (%SASA=33, pka = 16.00) | Y443($\%_{SASA}$=61, 0.42), substrate's COO- group (%SASA=92, 0.24) | | [+2] | -2.19 |
| | | | | D389($\%_{SASA}$=23, 0.44), E527($\%_{SASA}$=7, 2.68), Y526($\%_{SASA}$=57, 0.69), substrate's COO- group (%SASA=61, 1.17) | | [+1] | |
| | | | | D451($\%_{SASA}$=12, 0.42), Y629($\%_{SASA}$=0, 0.77), E535 (%SASA=0, 0.03), substrate's COO- group (%SASA=29, 0.26) | $Ca^{2+}$ (-1.25), R596(%SASA=0, -0.18) | [-1] | |

#### Table S4a

Table S4a: The distribution of catalytic base's pka determinants at active site of GlcA cleaving PLs belonging to family PL-5, and 8

| Protein name | PL-family | Optimum pH | Catalytic base | pka raising determinants | pka lowering determinants | Subsites | pka change due to desolvation of catalytic base |
|---|---|---|---|---|---|---|---|
| SpnHL (apo) | 8 | 6 | H399 (%SASA=17, pka= 7.91) | R336 (%SASA=72, -0.03) | E388 (%SASA=37, 0.29) | [+2] | -2.07 |
| | | | | R462 (%SASA=13, -0.55) | E577 (%SASA=14, 2.83), D398 (%SASA=0, 0.93) | [+1] | |
| SpnHL (substrate bound) | | | H399 (%SASA=2, pka=4.29) | R336 (%SASA=69, -0.02) | | [+2] | -3.21 |
| | | | | R462 (%SASA=0, -0.86) | E577 (%SASA=0, 1.72), substrate ring O atom (%SASA=4, 0.16) | [+1] | |
| CslA (apo) | 8 | 8 | H225 (%SASA=22, pka=6.83 | | E214 (%SASA=37, 0.32) | [+2] | -2.25 |
| | | | | R288 (%SASA=12, -0.56) | E371 (%SASA=10, 2.82) | [+1] | |
| CslA (substrate bound) | | | H225 (%SASA=8, pka=8.19 | R368 (%SASA=66, -0.01) | E214 (%SASA=20, 0.49) | [+2] | -3.17 |
| | | | | R288 (%SASA=0, -0.74) | E371 (%SASA=0, 3.62) | [+1] | |
| ArthroAC (apo) | 8 | 6 (more active for HA) | H233 (%SASA=37, pka=6.16) | | | [+2] | -1.22 |
| | | | | R296 (%SASA=30, -0.35) | H233 (0.09), D222 (%SASA=40, 0.16), E407 (%SASA=13, 0.99) | [+1] | |
| ArthroAC (CS-1) | | | H233 (%SASA=0, pka=9.71) | | Substrate's sulphate group (ASG, 0.21) | [+2] | |
| | | | | R296 (%SASA=0, -0.78) | D222 (%SASA=15, 0.65), E407 (%SASA=0, 3.57), substrate ring O atom (%SASA=0, 0.16), substrate's COO- group (%SASA=0, 3.03) | [+1] | |
| ArthroAC (CS-2) | | | H233 (%SASA=0, pka=6.41) | | | [+2] | -3.03 |
| | | | | R296 (%SASA=0, -0.69) | D222 (%SASA=15, 0.75), E407 (%SASA=0, 1.75), substrate's COOH group (%SASA=0, 1.13) | [+1] | |
| ArthroAC (HA) | | | H233 (%SASA=0, pka=6.38) | | | [+2] | |
| | | | | R296 (%SASA=0, -0.69) | D222 (%SASA=18, 0.69), E407 (%SASA=0, 1.75), substrate's COOH group (%SASA=0, 1.14) | [+1] | |

#### Table S4b

Table S4b: The distribution of catalytic base's pka determinants at active site of GlcA cleaving PLs belonging to family PL-5

| Protein name | PL-family | Optimum pH | Catalytic base | pka lowering determinants | pka raising determinants | Subsites | pka change due to desolvation of catalytic base |
|---|---|---|---|---|---|---|---|
| Smlt1473 (apo) | 5 | 5 | H168 (%SASA=0, pka=3.14) | R215 (%SASA=30, -0.36) | | [+1] | -3 |
| | | | R215 (%SASA=30, pka=12.72) | | D41 (%SASA=100, 0.09) | [+2] | -2.08 |
| | | | | | Y222 (%SASA=11, 1.23), Y39 (%SASA=38, 0.39), Y115 (%SASA=31, 0.05), E212 (%SASA=4, 0.49) | [+1] | |
| | | | | | Y38 (%SASA=33, 0.03), Y225 (%SASA=0, 0.01) | [-1] | |
| | | | Y222 (%SASA=11, pka=14.61) | | D41 (%SASA=100, 0.01) | [+2] | 2.56 |
| | | | | R215 (%SASA= , -1.23) | Y39 (%SASA=38, 0.99), Y115 (%SASA=31, 0.02), E212 (%SASA=4, 0.29) | [+1] | |
| | | | | R218 (%SASA= , -0.19) | Y38 (%SASA=33, 0.51), Y225 (%SASA=0, 0.79) | [-1] | |
| Smlt1473 (HA) | | | H168 (%SASA=0, pka=5.65) | | D111 (%SASA=55, 0.05) | [+2] | -3.95 |
| | | | | R215 (%SASA=12, -0.48) | substrate ring O atom (%SASA=0, 0.78), substrate's COO- group (%SASA=0, 1.99) , 1-->3 glycosidic Oxygen (%SASA=0, 0.75) | [+1] | |
| | | | R215 (%SASA=12, pka=14.20) | | D41 (%SASA=88, 0.11) | [+2] | -2.08 |
| | | | | | Y222 (%SASA=7, 1.51), Y39 (%SASA=28, 0.54), Y115 (%SASA=9, 0.11), E212 (%SASA=0, 0.84), substrate's COO- group (%SASA=0, 2.05) | [+1] | |
| | | | | | Y38 (%SASA=11, 0.22), Y225 (%SASA=0, 0.0) | [-1] | |
| | | | Y222 (%SASA=7, pka=17.70) | | D41 (%SASA=88, 0.09) | [+2] | 4.68 |
| | | | | R215 (%SASA=12 , -1.51), 1-->4 glycosidic Oxygen (%SASA=0, -0.07) | Y39 (%SASA=28, 2.03), E212 (%SASA=0, 0.27), substrate's COO- group (%SASA=0, 1.50) | [+1] | |
| | | | | R218 (%SASA=50, -0.45) | Y38 (%SASA=11, 0.96), Y225 (%SASA=0, 0.21) | [-1] | |

### Supplementary Information

#### Table S5

| Table S5: The distribution of catalytic base's pka determinants at active site of IdoA cleaving PLs belonging to family PL-13, and 24 | | | | | | | |
|---|---|---|---|---|---|---|---|
| **Protein name** | **PL-family** | **Optimum pH** | **Catalytic base** | **pka raising determinants** | **pka lowering determinants** | **Subsites** | **pka change due to desolvation of catalytic base** |
| **Heparinase-I (apo)** | 13 | 7.5 | H151 (%SASA=0, pka=1.71) | | R83(%SASA=6, -0.51) | [+2] | -2.73 |
| | | | | | K353(%SASA=0, -1.42) | [+1] | |
| | | | | | R344(%SASA=0, -0.13) | [-1] | |
| **Heparinase-I (substrate bound)** | | | H151 (%SASA=0, pka=3.09) | substrate's COO- group (%SASA=0, 0.15) | | [+3] | -3.25 |
| | | | | 1-->4 glycosidic Oxygen (%SASA=0, 0.36) | R83(%SASA=0, -0.39), substrate N2 (-0.39) | [+2] | |
| | | | | G152(%SASA=0, 0.49), substrate ring O atom (%SASA=0, 0.16), 1-->4 glycosidic Oxygen (%SASA=0, 0.36), substrate's COO- group (%SASA=0, 1.26) | K81(%SASA=0, -0.04), K252(%SASA=0, -0.02), R344(%SASA=0, -0.41) | [+1] | |
| | | | | | K353(%SASA=0, -1.11), substrate N2 (-0.22) | [-1] | |
| **LOR_107 (apo)** | 24 | 7.5 | H146(%SASA=18, pka=5.40) | E144(%SASA=3, 2.85) | H167(%SASA=11, -1.27), R259(%SASA=8, -0.32) | [+1] | -2.37 |
| **LOR_107 (substrate bound)** | | | H146(%SASA=0, pka=5.53) | E144(%SASA=3, 3.41), substrate ring O atom (%SASA=7, 0.21), 1-->4 glycosidic Oxygen (%SASA=7, 0.75), substrate's COO- group (%SASA=7, 1.32) | H167(%SASA=11, -2.03), R259(%SASA=8, -0.47) | [+1] | -4.19 |

#### Table S6

| Table S6: The distribution of catalytic base's pka determinants at active site of ManA cleaving PLs belonging to family PL-5 | | | | | | | |
|---|---|---|---|---|---|---|---|
| **Protein name** | **PL-family** | **Optimum pH** | **Catalytic base** | **pka raising determinants** | **pka lowering determinants** | **Subsites** | **pka change due to desolvation of catalytic base** |
| **Alginate Lyase A1-III (apo)** | 5 | 8 | H192 (%SASA=10, pka=6.89) | | R196(%SASA=18, -0.31) | [+2] | -2.71 |
| | | | | E140 (%SASA=0, 3.34), E236(%SASA=5, 0.23) | R239 (%SASA=53, -0.17) | [+1] | |
| | | | R239 (%SASA=53, pka=12.54) | Y137(%SASA=55, 0.05), Y246 (%SASA=38, 0.67), E236 (%SASA=5, 0.36), E241 (%SASA=100, 0.02) | | [+1] | -1.04 |
| | | | Y246 (%SASA=38, pka=10.49) | E236(%SASA=5, 0.17), E241(%SASA=100, 0) | R239(%SASA=53, --0.67), | [+1] | 1.14 |
| | | | | | | [-1] | |
| | | | | D304(%SASA=100, 0.01), D314(%SASA=32, 0.03) | R306 (%SASA=85, -0.18) | [-2] | |
| **Alginate Lyase A1-III (substrate bound)** | | | H192 (%SASA=0, pka=8.22) | | R196(%SASA=18, -0.50) | [+2] | -3.94 |
| | | | | E140 (%SASA=0, 1.60), E236(%SASA=5, 0.30), substrate's COO- group (%SASA=7, 2.03), substrate ring O atom (%SASA=0, 0.72) | R239 (%SASA=53, -0.42) | [+1] | |
| | | | | substrate's COO- group (%SASA=7, 0.15) | | [-1] | |
| | | | R239 (%SASA=7, pka=15.20) | Y137(%SASA=55, 0.10), Y246 (%SASA=38, 1.95), E236 (%SASA=5, 0.84), E241 (%SASA=100, 0.05), Y68(%SASA=12, 0.76), substrate's COO- group (%SASA=0, 2.43) | | [+1] | -3.39 |
| | | | | | R67(%SASA=17, -0.04) | [-1] | |
| | | | Y246 (%SASA=0, pka=15.97) | E236(%SASA=5, 0.35), E241(%SASA=100, 0.12), Y68(%SASA=12, 2.6), substrate's COO- group (%SASA=7, 1.78) | R239(%SASA=53, -1.95), 1-->4 glycosidic Oxygen (%SASA=0, -0.28) | [+1] | 4.51 |
| | | | | Y80(%SASA=1, 0.05), substrate's COO- group (%SASA=7, 0.23) | R67(%SASA=17, -0.68), substrate C3-OH group (%SASA=0, -0.46) | [-1] | |
| | | | | D304(%SASA=100, 0.03), substrate's COO- group (%SASA=7, 0.01) | R306 (%SASA=85, -0.32) | [-2] | |
| **Smlt1473 (apo)** | 5 | 9 | H168(%SASA=0, pka=2.97) | | R215 (%SASA=27, -0.41) | [+1] | -3.09 |
| | | | | | R312(%SASA=0, -0.03) | [-1] | |
| | | | R215(%SASA=27, pka=12.90) | E212(%SASA=2 , 0.54), D41(%SASA=100, 0.06) | | [+2] | -2.33 |
| | | | | Y39(%SASA=37, 0.58), Y222 (%SASA=9, 1.33), Y115 (%SASA=30 , 0.10) | | [+1] | |
| | | | | Y38(%SASA=31 , 0.07), Y225(%SASA=0, 0.05) | | [-1] | |
| | | | Y222(%SASA=9, 14.79) | E212(%SASA=2 , 0.30), D41(%SASA=100, 0.06) | | [+2] | 2.82 |
| | | | | Y39 (%SASA=37 , 1.88), Y115(%SASA=30, 0.06) | R215 (%SASA=27, -1.33), R218(%SASA=69, -0.22) | [+1] | |
| | | | | Y38(%SASA=31 , 0.58), Y225(%SASA=0, 0.73) | R312(%SASA=0 , -0.02) | [-1] | |
| **Smlt1473 (substrate bound)** | | | H168(%SASA=0, pka=2.74) | substrate's COO- group (%SASA=0, 0.22), substrate ring O atom (%SASA=0, 0.61), D111(%SASA=59 , 0.08) | R215 (%SASA=18, -0.45) | [+1] | -4.23 |
| | | | | | | [-1] | |
| | | | R215(%SASA=18, pka=14.98) | E212(%SASA=0 , 0.54), D41(%SASA=93 , 0.06) | | [+2] | -2.33 |
| | | | | Y39(%SASA=27 , 0.58), Y222 (%SASA=0, 1.33), Y115 (%SASA=17 , 0.10) | | [+1] | |
| | | | | Y38(%SASA=9 , 0.07), Y225(%SASA=0 , 0.05) | | [-1] | |
| | | | Y222(%SASA=0, 17.99) | E212(%SASA=0 , 0.31), D41(%SASA=93 , 0.02) | | [+2] | 4.8 |
| | | | | Y39 (%SASA=27 , 2.26), Y115(%SASA=17, 0.06), substrate's COO- group (%SASA=0, 1.71) | R215 (%SASA=18, -1.52), R218(%SASA=61 , -0.27), 1-->4 glycosidic Oxygen (%SASA=0, -0.85), substrate C3-OH group (%SASA=0, -0.85) | [+1] | |
| | | | | Y38(%SASA=9 , 1.23), Y225(%SASA=0 , 0.53), substrate's COO- group (%SASA=0, 0.53) | R312(%SASA=0 , -0.02) | [-1] | |
| | | | | substrate's COO- group (%SASA=45, 0.10) | | [-2] | |

#### Table S7

Table S7: The distribution of catalytic base's pka determinants at active site of GulA cleaving PLs belonging to family PL-6, and 7

| Protein name | PL-family | Optimum pH | Catalytic base | pka raising determinants | pka lowering determinants | Subsites | pka change due to desolvation of catalytic base |
|---|---|---|---|---|---|---|---|
| AlyF (apo) | 6 | 7.5 | K272(%SASA=0, pka=9.34) | D242(%SASA=43, 0.20) | R239(%SASA=0,-0.75) | [+3] | -2.13 |
| | | | | | | [+2] | |
| | | | | E265(%SASA=0, 0.40), E236(%SASA=0, 0.99), E267(%SASA=0, 1.24) | | [+1] | |
| | | | | | R266(%SASA=0, -0.36), R293(%SASA=0, -0.76) | [-1] | |
| AlyF (substrate bound) | | | K272(%SASA=0, pka=10.95) | D242(%SASA=26, 0.85), substrate's COO- group (%SASA=11, 0.45) | R239(%SASA=0,-1.09) | [+3] | -4.87 |
| | | | | substrate's COO- group (%SASA=20, 2.88) | | [+2] | |
| | | | | E265(%SASA=0, 0.10), E236(%SASA=0, 0.71), E267(%SASA=0, 0.59), substrate's COO- group (%SASA=0, 1.29) | | [+1] | |
| | | | | substrate's COO- group (%SASA=0, 0.06) | R266(%SASA=0, -0.05), R293(%SASA=0, -0.49) | [-1] | |
| AlyF (substrate bound, Ca2+ modelled) | | | K272(%SASA=0, pka=9.46) | D242(%SASA=32, 0.32), substrate's COO- group (%SASA=31, 0.27) | K165(%SASA=33, -0.01), R239(%SASA=0,-1.09) | [+3] | -4.25 |
| | | | | substrate's COO- group (%SASA=30, 1.34) | | [+2] | |
| | | | | E265(%SASA=0, 0.52), E236(%SASA=0,1.42), E267(%SASA=0,1.10), substrate's COO- group (%SASA=0, 2.24) | Ca2+ (-2.59) | [+1] | |
| | | | | substrate's COO- group (%SASA=0, 0.19) | R293(%SASA=0, -0.59) | [-1] | |
| AlyGC (apo) | 6 | 7 | K220(%SASA=24, pka=9.59) | | R187(%SASA=13, -0.52) | [+3] | -1.59 |
| | | | | E184(%SASA=13, 0.57) | | [+2] | |
| | | | | E213(%SASA=17, 0.20), E215(%SASA=0, 0.81) | Ca2+ (-1.95), R241(%SASA=22, -0.38) | [+1] | |
| | | | | | | [-1] | |
| AlyGC (GulA bound) | | | K220(%SASA=9, pka=6.46) | substrate's COO- group (%SASA=53, 0.10) | R187(%SASA=0, -0.74) | [+3] | -4.89 |
| | | | | E184(%SASA=8, 0.67), substrate's COO- group (%SASA=65, 0.57) | | [+2] | |
| | | | | E213(%SASA=11, 0.25), E215(%SASA=0, 1.28), substrate's COO- group (%SASA=26, 2.02) | Ca2+ (-2.86), R241(%SASA=15, -0.56) | [+1] | |
| | | | | substrate's COO- group (%SASA=19, 0.22) | R265(%SASA=0, -0.02) | [-1] | |
| AlyGC (ManA bound) | | | K220(%SASA=0, pka=7.23) | | R187(%SASA=, -0.83) | [+3] | -4.49 |
| | | | | E184(%SASA=, 0.91), substrate's COO- group (%SASA=, 0.18) | | [+2] | |
| | | | | E213(%SASA=, 0.35), E215(%SASA=0, 1.28), substrate's COO- group (%SASA=, 2.23) | Ca2+ (-2.09), R241(%SASA=, -0.81) | [+1] | |
| | | | | | R265(%SASA=, -0.02) | [-1] | |
| Alginate lyase A1-II (apo) | 7 | 7.5 | H191(%SASA=0, pka=3.54) | | R150(%SASA=24, -0.16) | [+3] | -2.74 |
| | | | | | K280(%SASA=0, -0.85) | [+2] | |
| | | | | G192's backbone O (0.83), E148(%SASA=0, 0.37) | R146(%SASA=10, -0.37) | [+1] | |
| | | | | | K205(%SASA=35, -0.04) | [-1] | |
| Alginate lyase A1-II (GulA bound) | | | H191(%SASA=0, pka=5.83) | substrate's COO- group (%SASA=31, 0.40) | R150(%SASA=12, -0.22) | [+3] | -3.46 |
| | | | | substrate's COO- group (%SASA=75, 0.10) | K280(%SASA=0, -1.12) | [+2] | |
| | | | | E148(%SASA=, 0.82), substrate's COO- group (%SASA=16, 3.63) | R146(%SASA=0, -0.69) | [+1] | |
| | | | | substrate's COO- group (%SASA=69, 0.02) | K205(%SASA=22, -0.15) | [-1] | |

**A**

| Substrate | PL Family | Structural Folds |
|---|---|---|
| GalA | PL-1, 2, 3, 9, 10, 11, 22, 26 | Parallel β-helix [PL-1, 3, 9], $(\alpha/\alpha)_7$ barrel [PL-2], $(\alpha/\alpha)_3$ barrel [PL-10], $(\alpha/\alpha)_6$ barrel [PL-26], β-propeller [PL-11, 22] |
| GlcA/IdoA | PL-5, 6, 7, 8, 12, 13, 14, 15, 20, 21, 23, 24, 25, 28, 31 | $(\alpha/\alpha)_6$ barrel [PL-5], parallel β-helix [PL-6, 31], β-jellyroll [PL-7, 13, 14, 20, 28], $(\alpha/\alpha)_6$ barrel + anti-parallel β-sheet [PL-8, 15, 21], $(\alpha/\alpha)_5$ barrel + anti-parallel β-sheet [PL-12, 23], β-propeller [PL-24, 25] |
| ManA/GulA | PL-5, 6, 7, 14, 15, 17, 18, 31, 39 | β-jellyroll [PL-18], $(\alpha/\alpha)_6$ barrel + anti-parallel β-sheet [PL-17, 39] |

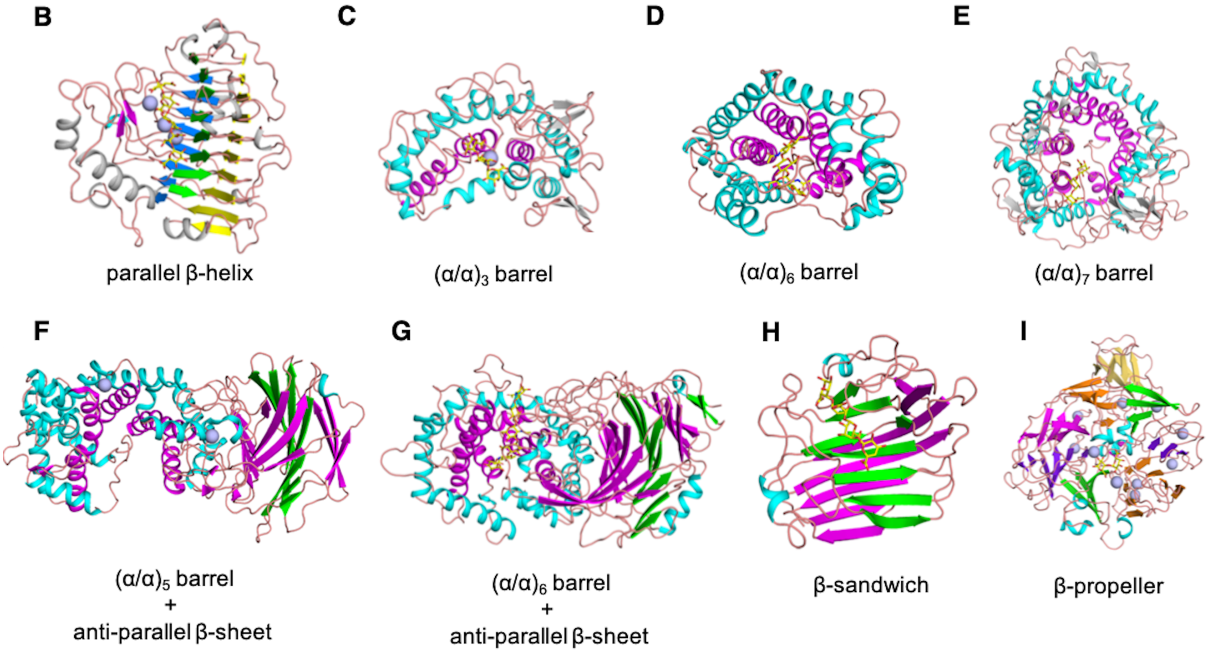

**FIGURE S1 Structural diversity among PLs grouped as a function of substrate.** (A) Table showing PL families and their respective protein fold grouped according to the major substrate type PLs bind at [+1] subsite, GalA, GlcA/IdoA, ManA/GulA. (B-I) Cartoon representation of various folds observed among PL families (A), the different colours have been used to highlight the structural folds.

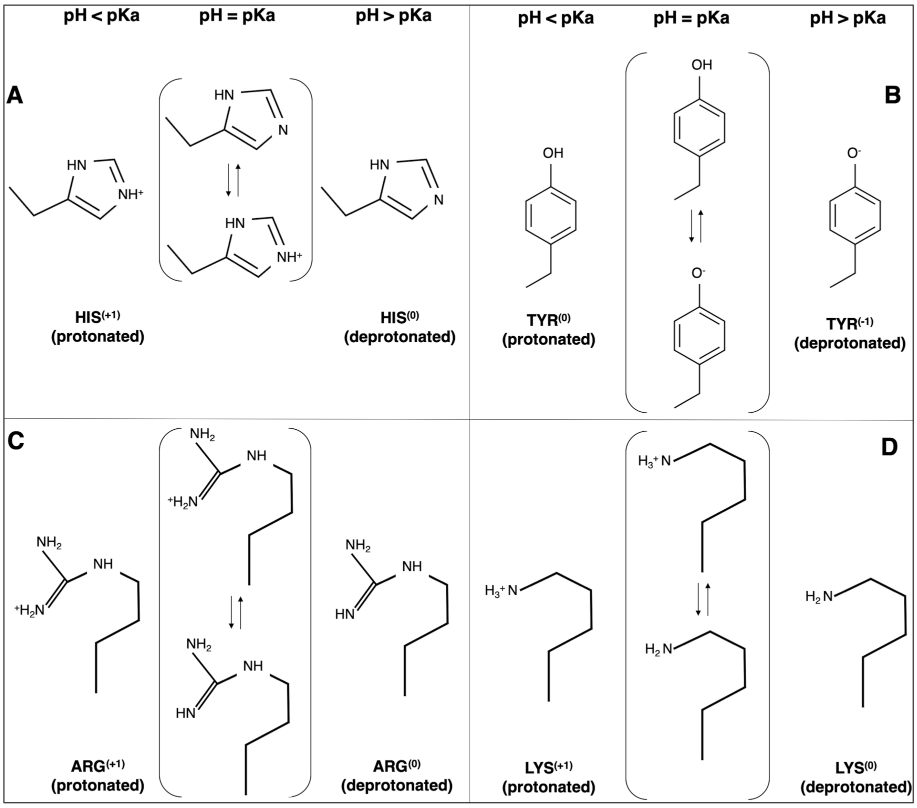

**FIGURE S2 Schematics showing the charge state (pH *vs* p$K_a$) of side chain of titrable amino acids implicated as catalytic base among PLs.** The basic amino acids such as His (imidazolium), Arg (guanidine), and Lys (butylammonium) side chain remain protonated (positively charged) at pH below their p$K_a$ , and get deprotonated (neutral) at pH above their p$K_a$. Only in deprotonated state, His, Arg, and Lys can act as catalytic base; and likewise, Tyr (hydroxyl) side chain remains protonated (uncharged) at pH below its p$K_a$. Whereas, the pH above its p$K_a$ it loses the proton from its side-chain to acquire negative charge, and become catalytic base candidate. For all titrable amino acids, the p$K_a$ is defined as pH at which the amount of protonated and deprotonated states are at equilibrium. The p$K_a$ is a relative value, and for amino acids constituting protein, its value depend on many intrinsic and extrinsic factors (please refer to Note S1 for more details).

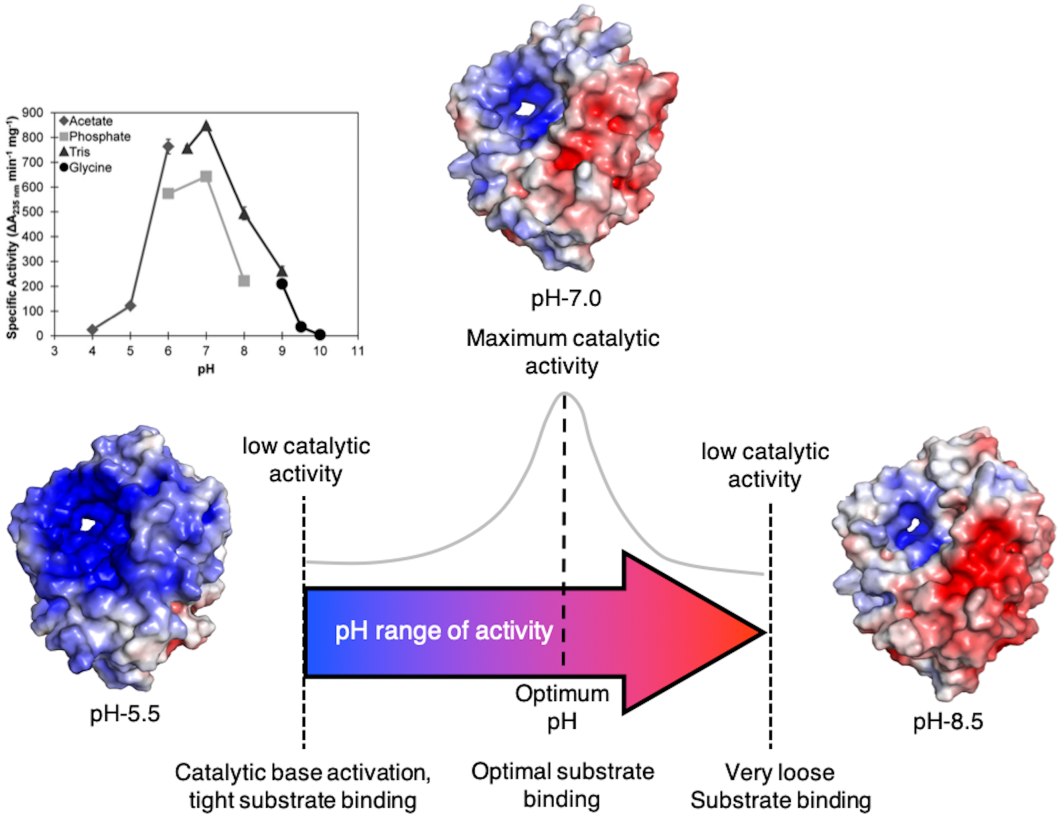

**FIGURE S3 The correlation of electrostatic potential surface charge and Smlt1473 specific activity against poly-Glucuronate as a function of pH.** The specific activity ($\Delta A_{235\ nm}$ $min^{-1}$ $mg^{-1}$) measured as a function of pH shows Smlt1473's pH range of activity toward poly-Glucuronate from pH ~5.0-9.0, the maximum activity is observed at pH-7.0. The electrostatic potential surface charge (-5 to +5 $K_bT/e_c$) at pH 5.5, 7.0, and 8.5 shows a loss in electropositive surface around the substrate binding catalytic tunnel with increase in pH. At higher pH the electropositive surface confined only toward the active core of catalytic tunnel. The schematics of our rationale of PL-pH relationship propose that, at low pH the negatively charged anionic polysaccharide substrate will bind very tightly with highly electropositive catalytic tunnel resulting into negligible catalytic activity; and also at such low pH, the proposed catalytic base will be protonated i.e. $pk_a > pH$ (no activity). As the pH rise, at certain pH ($pk_a \approx pH$) the catalytic base will activate (deprotonation) and start acting as base, but still the high precedence of electropositive surface charge (tight substrate binding) will result in low activity. However, if pH continue rising ($pk_a > pH$) the activity will also increase due to decrease in substrate binding (due to decreasing PL's electropositive surface charge); and reach a pH where the maximal activity will be observed [$pH_{(optimum\ activity)}$ or $pH_{(optimal\ substrate\ binding)}$]. After further increase in pH, the activity start declining due to decrease in electropositive surface charge (concomitant increase in electronegative charge surface) resulting in very low substrate binding. Finally, at above and last point of catalytic pH range, the enzymatic activity diminished due to overwhelming negative surface charge which result into unfavourable interaction with negatively charged anionic polysaccharide substrate. This hypothesis of pH and catalytic activity relationship hold only true for enzymes like PL which act on charged substrate as well utilize titrable amino acids for catalysis.

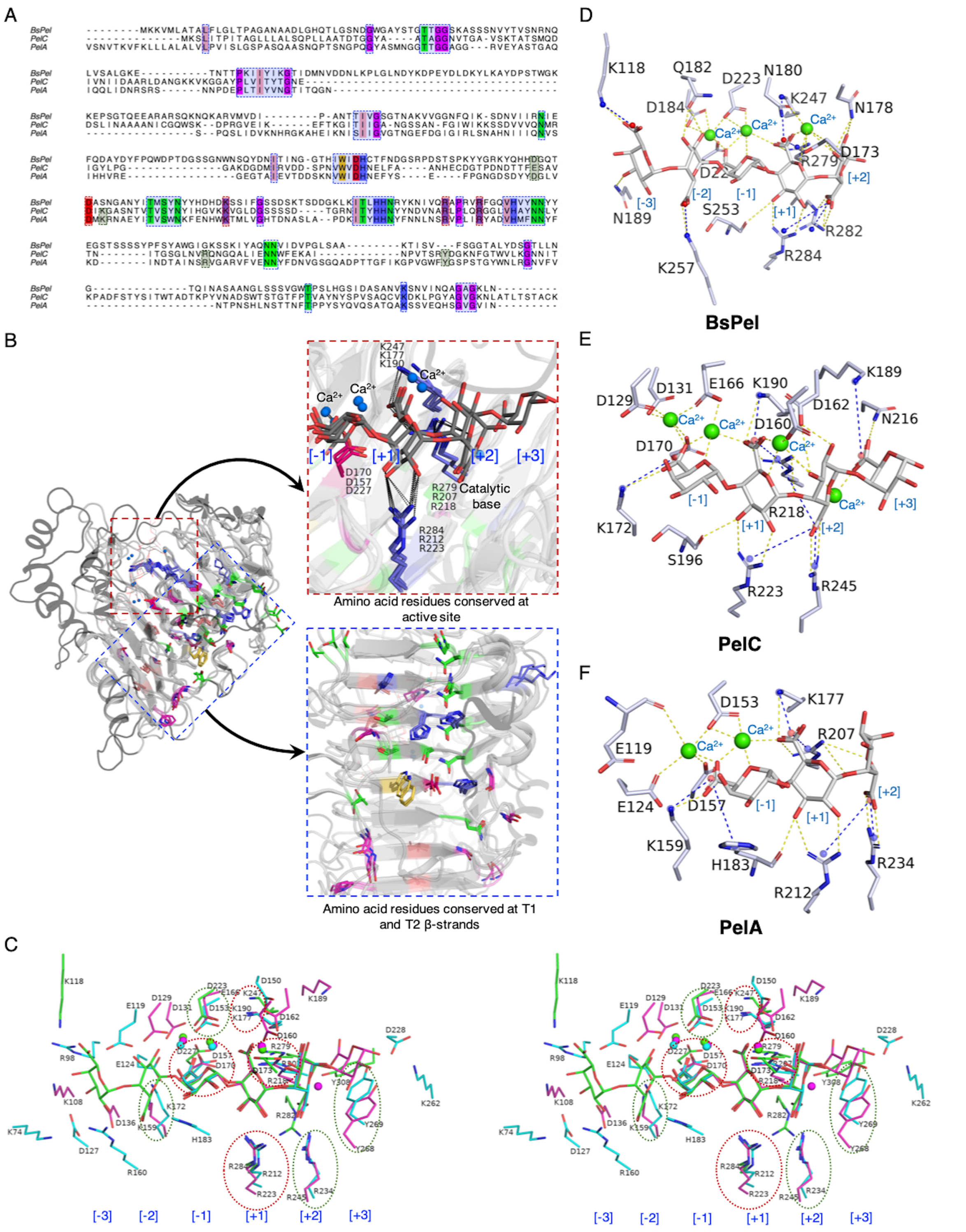

**FIGURE S4. Sequence-Structure comparison of PL-1 family of pectate lyases**. **(A)** Structure guided sequence alignment of BsPel, PelC, and PelA, the conserved residues are highlighted by amino acid color code (salmon = aliphatic amino acids such as Ile, Leu, and Ala, green = polar uncharged amino acids like Asn, Gln, Thr, and Ser, purple = for Gly, and Pro, red = acidic amino acid such as Asp, and Glu, blue = basic amino acids such as His, Arg, and Lys, yellow = aromatic amino acids such as Trp, Tyr, and Phe). The conserved structural, and active site regions are displayed under blue and red dashed and shaded box respectively. **(B)** Structural alignment of BsPel,

PelC, and PelA is displayed as cartoon, the conserved amino acids are displayed as sticks which are color coded according to type of amino acids described in A, the region of sequence insertion cartoon model are colored black. The image inset in red and blue dashed box represent the conserved active site and structural region described in sequence alignment shown in A. **(C)** Stereographic image of titrable amino acids around the structurally aligned substrate complexed with BsPel (green stick), PelC (purple), and PelA (cyan). The amino acids conserved for all three selected PLs are shown in red dotted ovals, or otherwise green dotted oval for at least two PLs. **(D-F)** Enzyme-Substrate interaction derived from modelled (catalytic base is computationally mutated back to wildtype) and relaxed (to remove steric clashes) substrate (GalA) complexed structure of selected PL-1 enzymes.

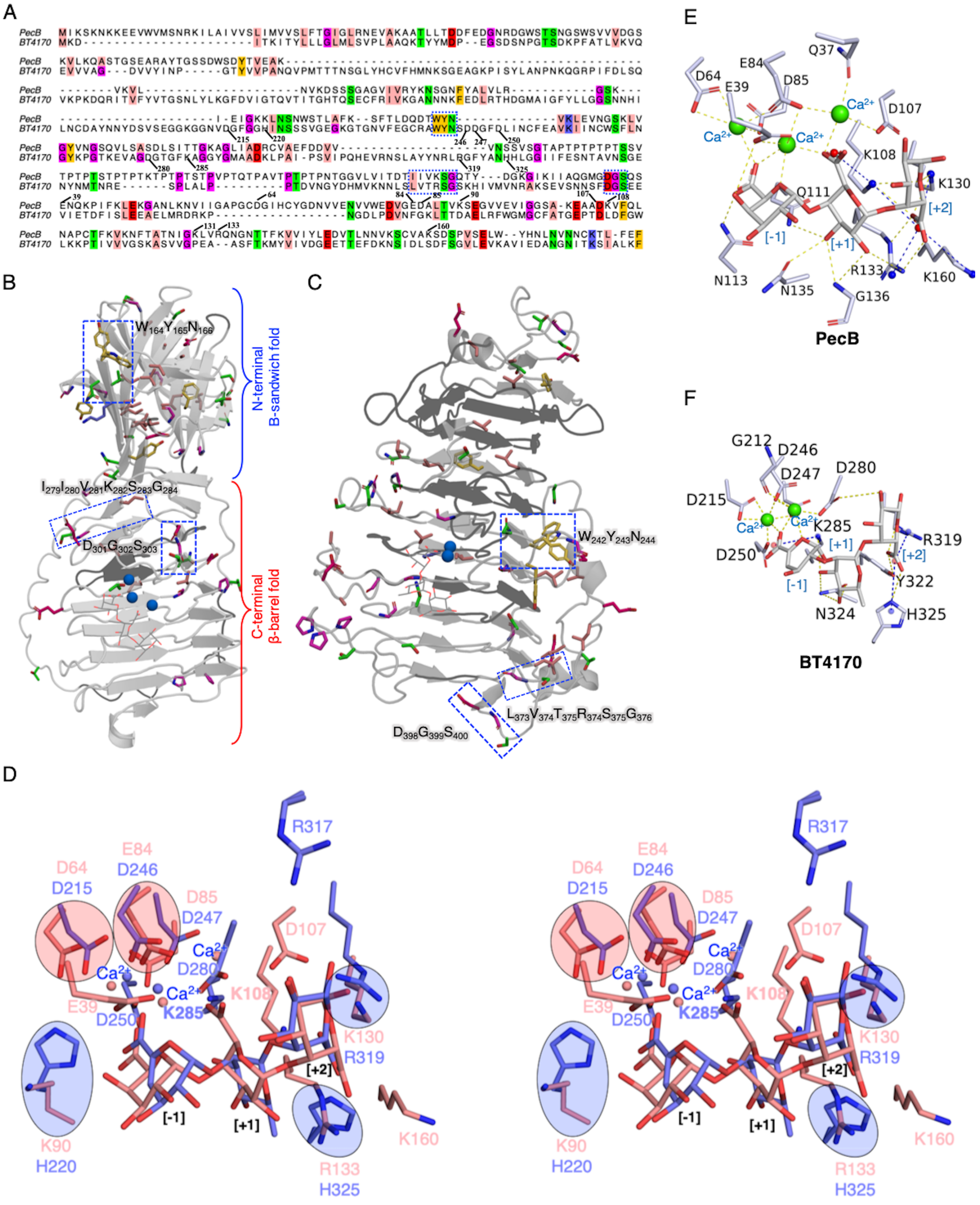

**FIGURE S5. Sequence-Structure comparison of PL-3, and 9 family of pectate lyases**. **(A)** Structure guided sequence alignment of PecB and BT4170, the conserved residues are highlighted by amino acid color code as discussed in Figure S4. The conserved structural regions are displayed under blue dashed and shaded box, the active sites are not sequentially conserved. **(B)** PecB, **(C)** BT4170 are displayed as cartoon model, the conserved amino acids are displayed as sticks which are color coded according to type of amino acids described in A, the region of sequence insertion cartoon model are colored black. The blue dashed box represent the conserved structural region described in sequence alignment shown in A. **(D)** Stereographic image of titrable amino acids around the structurally aligned substrate complexed with PecB (salmon stick), BT4170 (blue stick). The amino acids conserved for both PLs are shown in red (acidic amino acids) and blue (basic amino acids) shaded ovals. **(E-F)** Enzyme-Substrate interaction derived from modelled (catalytic base is computationally mutated back to wildtype) and relaxed (to remove steric clashes) substrate (GalA) complexed structure of selected PL-3, and 9 enzymes.

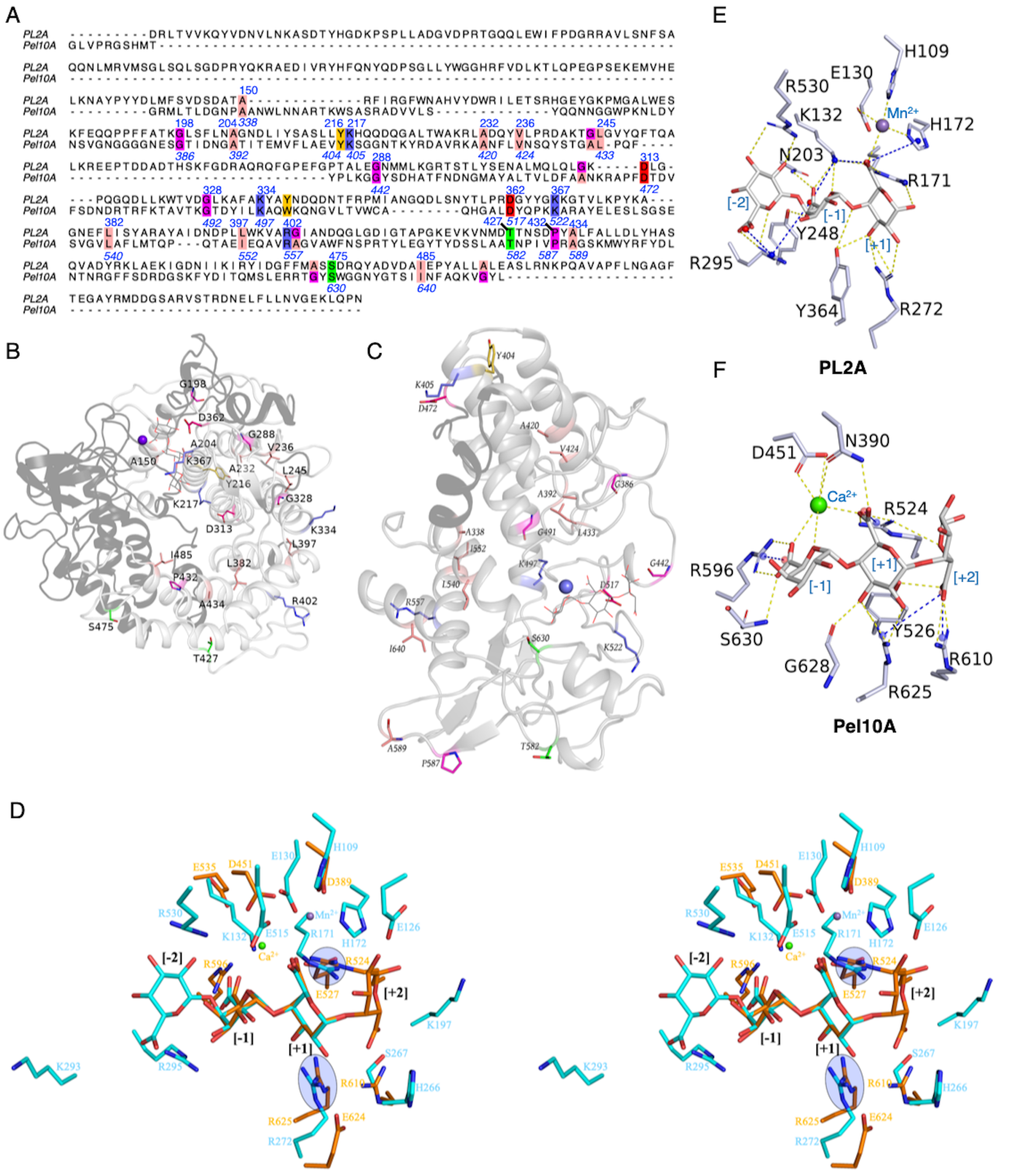

**FIGURE S6. Sequence-Structure comparison of PL-2, and 10 family of pectate lyases**. **(A)** Structure guided sequence alignment of PL2A and Pel10A, the conserved residues are highlighted by amino acid color code as discussed in Figure S4. The conserved structural regions are displayed under blue dashed and shaded box, but there is lack of continuous conserved sequence. **(B)** PL2A, **(C)** Pel10A are displayed as cartoon model, the conserved amino acids are displayed as sticks which are color coded according to type of amino acids described in A, the region of sequence insertion cartoon model are colored black. **(D)** Stereographic image of titrable amino acids around the structurally aligned substrate complexed with PL2A (cyan stick), Pel10A (dark orange stick). The amino acids conserved structurally for both PLs are shown in blue shaded ovals. **(E-F)** Enzyme-Substrate interaction derived from modelled (catalytic base is computationally mutated back to wildtype) and relaxed (to remove steric clashes) substrate (GalA) complexed structure of selected PL-2, and 10 enzymes.

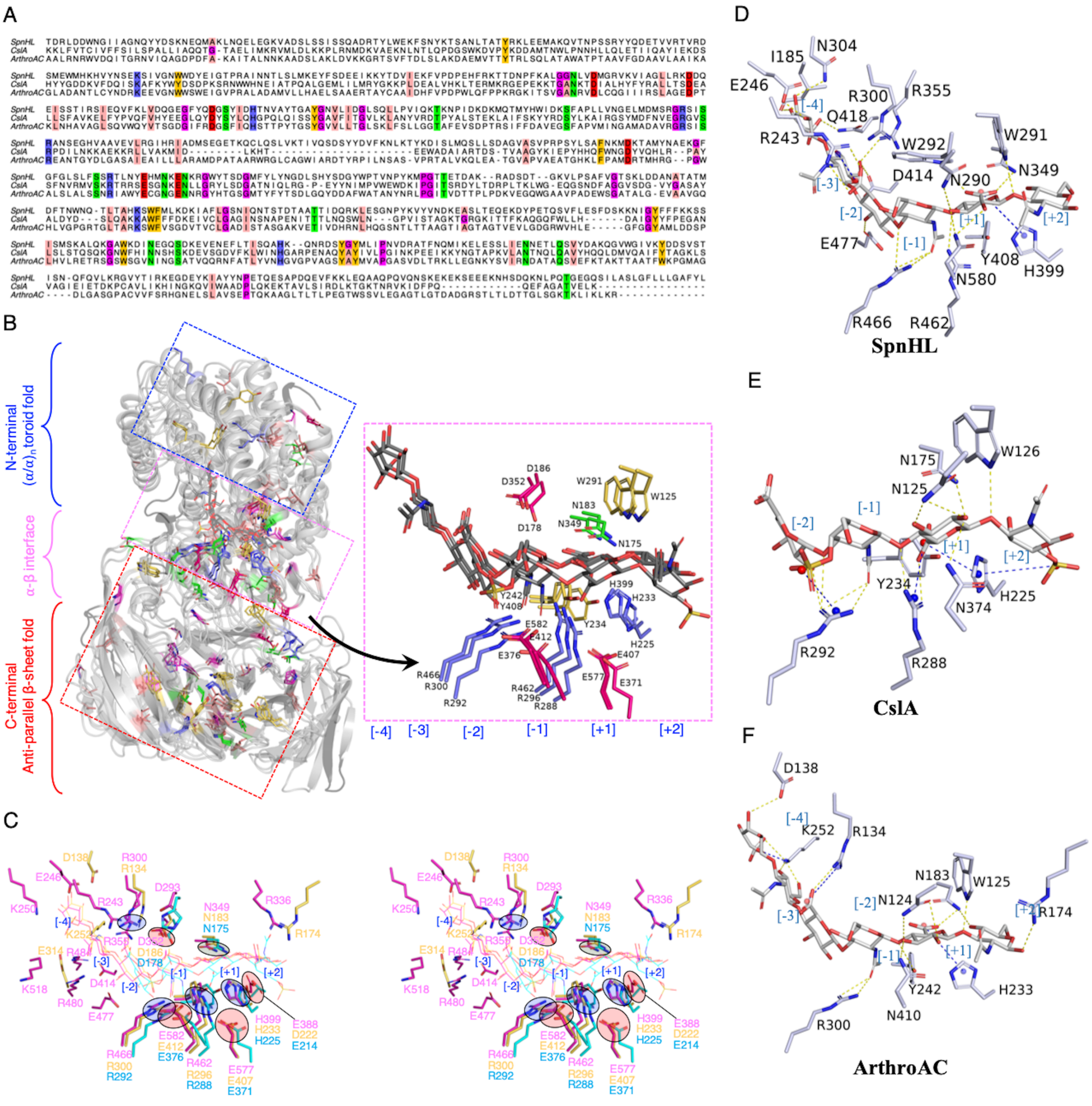

**FIGURE S7. Sequence-Structure comparison of PL-8 family of polyGlcA lyases**. **(A)** Structure guided sequence alignment of SpnHL (hyaluronate specific), CslA (chondroitin sulphate specific) and ArthroAC (bi-functional but more activity toward hyaluronic acid), the conserved residues are highlighted by amino acid color code as discussed in Figure S4. **(B)** Structural alignment of SpnHL, CslA, and ArthroAC is displayed as cartoon, the conserved amino acids are displayed as sticks which are color coded according to type of amino acids described in A, the region of sequence insertion cartoon model are colored black. The structural section marked in blue, magenta and red dashed box contains the conserved amino acid in N-terminal $(\alpha/\alpha)_n$ toroid fold, $\alpha$-$\beta$ interface (active site), and C-terminal anti-parallel $\beta$-sheet region of selected PL-8 enzymes. The image inset zoomed the aligned active site showing conserved amino acids common to the selected PLs. **(C)** Stereographic image of titrable amino acids around the structurally aligned substrate complexed with SpnHL (magenta stick), CslA (cyan stick), and ArthroAC (yellow stick). The acidic and basic amino acids conserved for all PLs are shown in red, and blue shaded ovals respectively. **(D-F)** Enzyme-Substrate interaction derived from modelled (catalytic base is computationally mutated back to wildtype) and relaxed (to remove steric clashes) substrate (GalA) complexed structure of selected PL-8 enzymes.

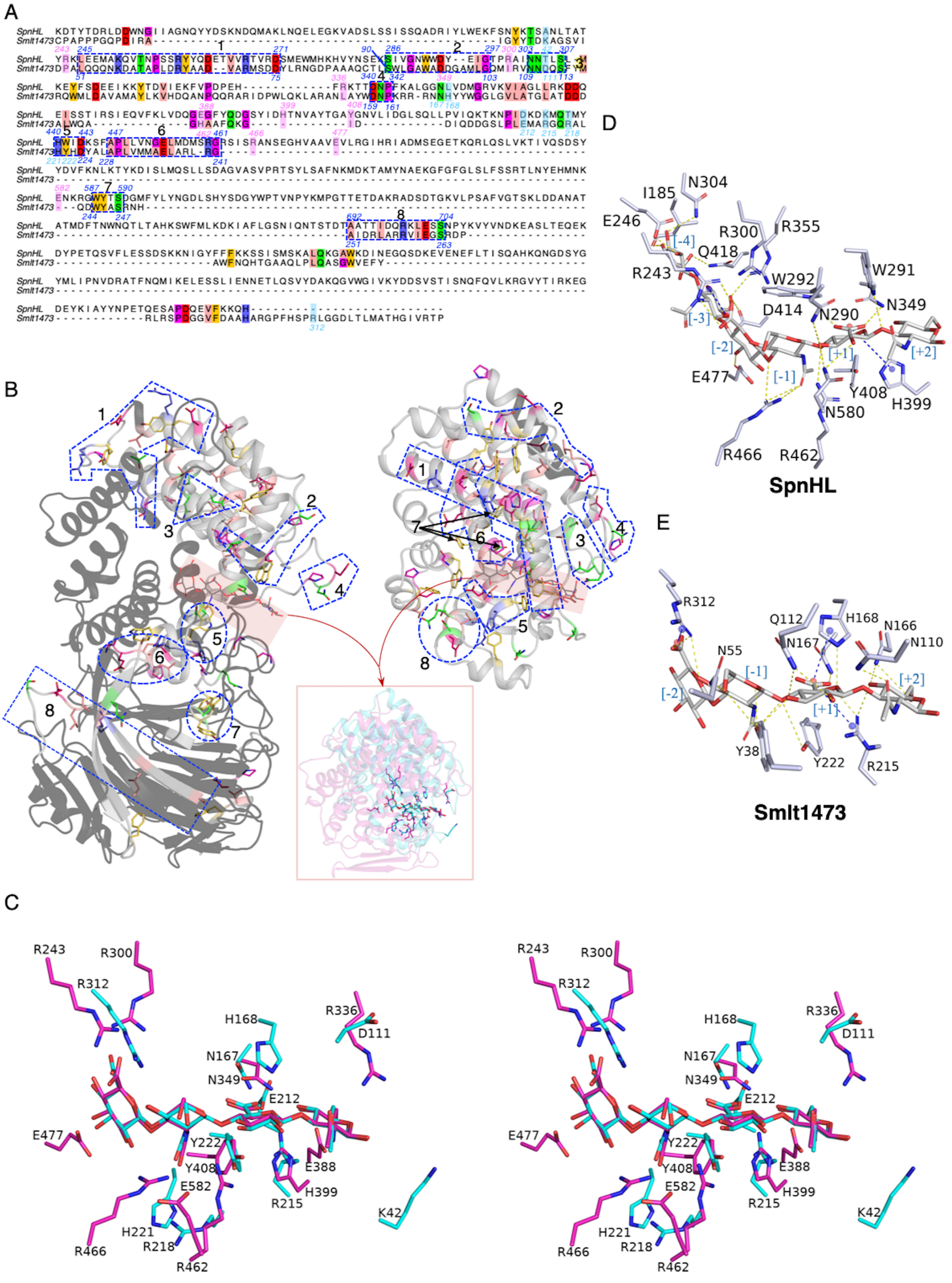

**FIGURE S8. Sequence-Structure comparison of PL-5 and PL-8 family of polyGlcA lyases**. **(A)** Structure guided sequence alignment of SpnHL (hyaluronate specific), and Smlt1473 (multi-functional PL) the conserved residues are highlighted by amino acid color code as discussed in Figure S4. **(B)** Structural alignment of SpnHL, and Smlt1473 is displayed as cartoon, the conserved amino acids are displayed as sticks which are color coded according to type of amino acids described in A, the region of sequence insertion cartoon model are colored black. The conserved structural region based on sequence alignment are marked under blue dashed box. The image inset zooms the aligned active site showing conserved amino acids common to the selected PLs. **(C)** Stereographic image of titrable amino acids around the structurally aligned substrate complexed with SpnHL (magenta stick), and Smlt1473 (cyan stick). **(D-F)** Enzyme-Substrate interaction derived from modelled (catalytic base is computationally mutated back to wildtype) and relaxed (to remove steric clashes) substrate (GalA) complexed structure of selected PL-5 and 8 enzymes.

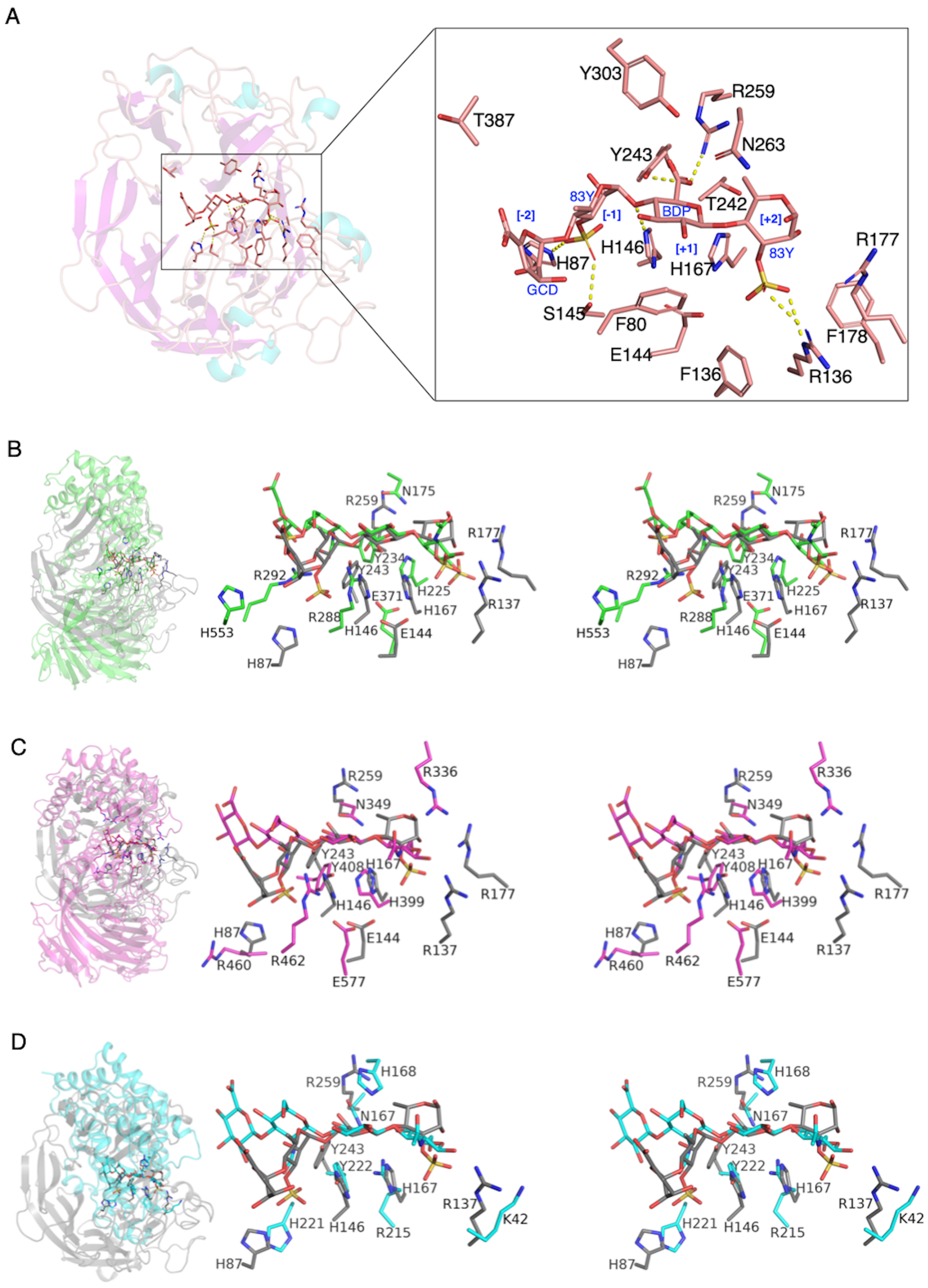

**FIGURE S9. Structural comparison of PL (LOR_107, PL-24) cleaving ulvan (non-glycan GlcA containing substrate) with PLs cleaving GlcA containing glycan substrate CslA (PL-8), SpnHL (PL-8), and Smlt1473 (PL-5).**

Supplementary Information

**(A)** Structural description of LOR_107 active site, inset zooms the active site enzyme-substrate interaction. Substrate coordinate based structural alignment of LOR_107 with **(B)** CslA which specifically cleaves chondroitin sulphate at pH-8.0, **(C)** SpnHL which specifically cleaves hyaluronic acid at pH-6.0, and **(D)** Smlt1473 a multifunctional PL which cleaves hyaluronate, glucuronate, and mannuronate at pH-5.0, 7.0, and 9.0.

Supplementary Information

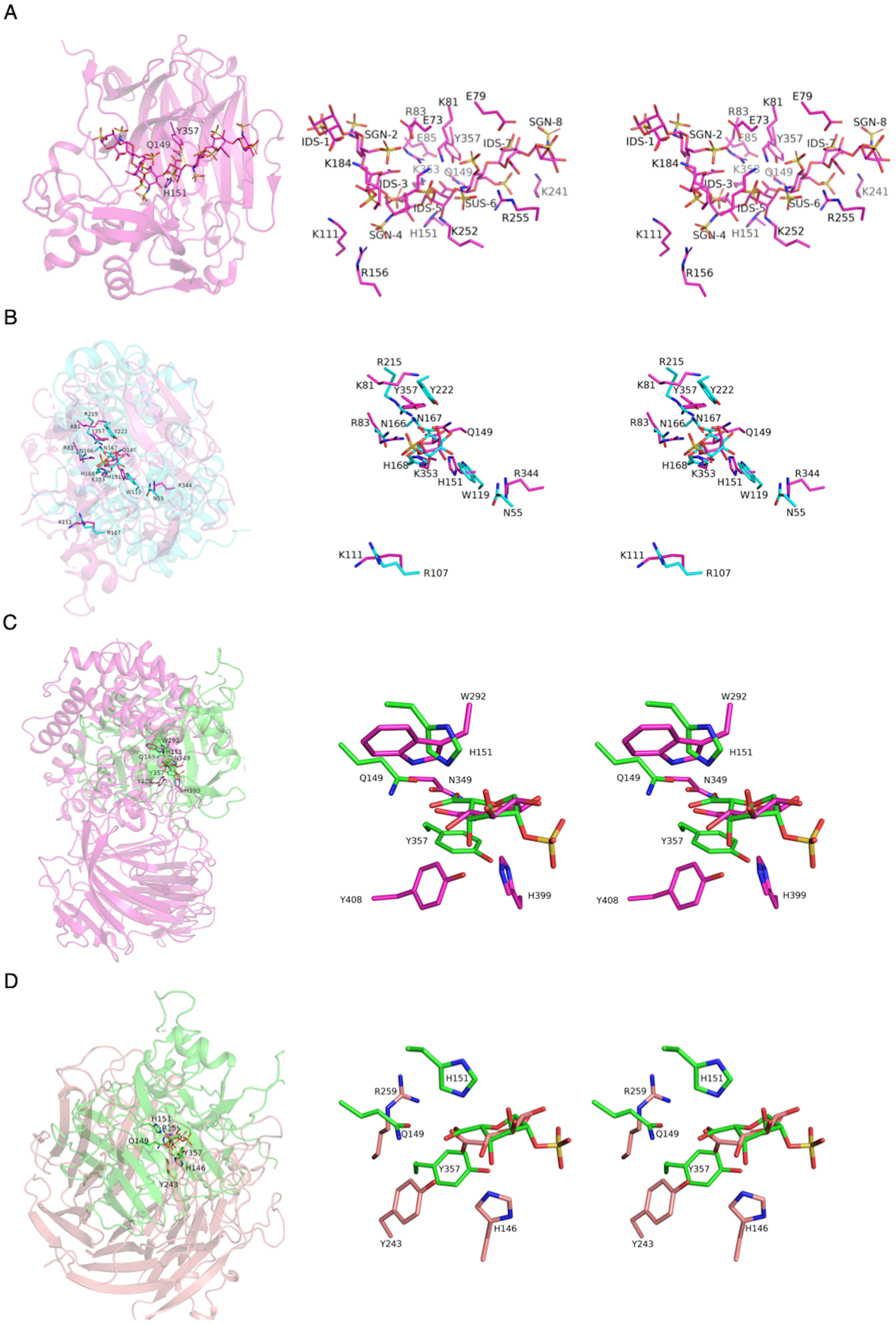

**FIGURE S10. Structural comparison of PL (Heparinase-I, PL-13) cleaving Heparin sulphate (IdoA containing substrate) with PLs cleaving GlcA containing substrate, Smlt1473 (PL-5), SpnHL (PL-8), and LOR_107 (PL-24).** **(A)** Structural description of Heparinase-I active site, and stereo image of the active site. Substrate coordinate based structural alignment of Heparinase-I with **(B)** Smlt1473 a multifunctional PL which cleaves hyaluronate, glucuronate, and mannuronate at pH-5.0, 7.0, and 9.0 **(C)** SpnHL which specifically cleaves hyaluronic acid at pH-6.0, and **(D)** LOR_107 which cleaves ulvan at pH-7.5.

Supplementary Information

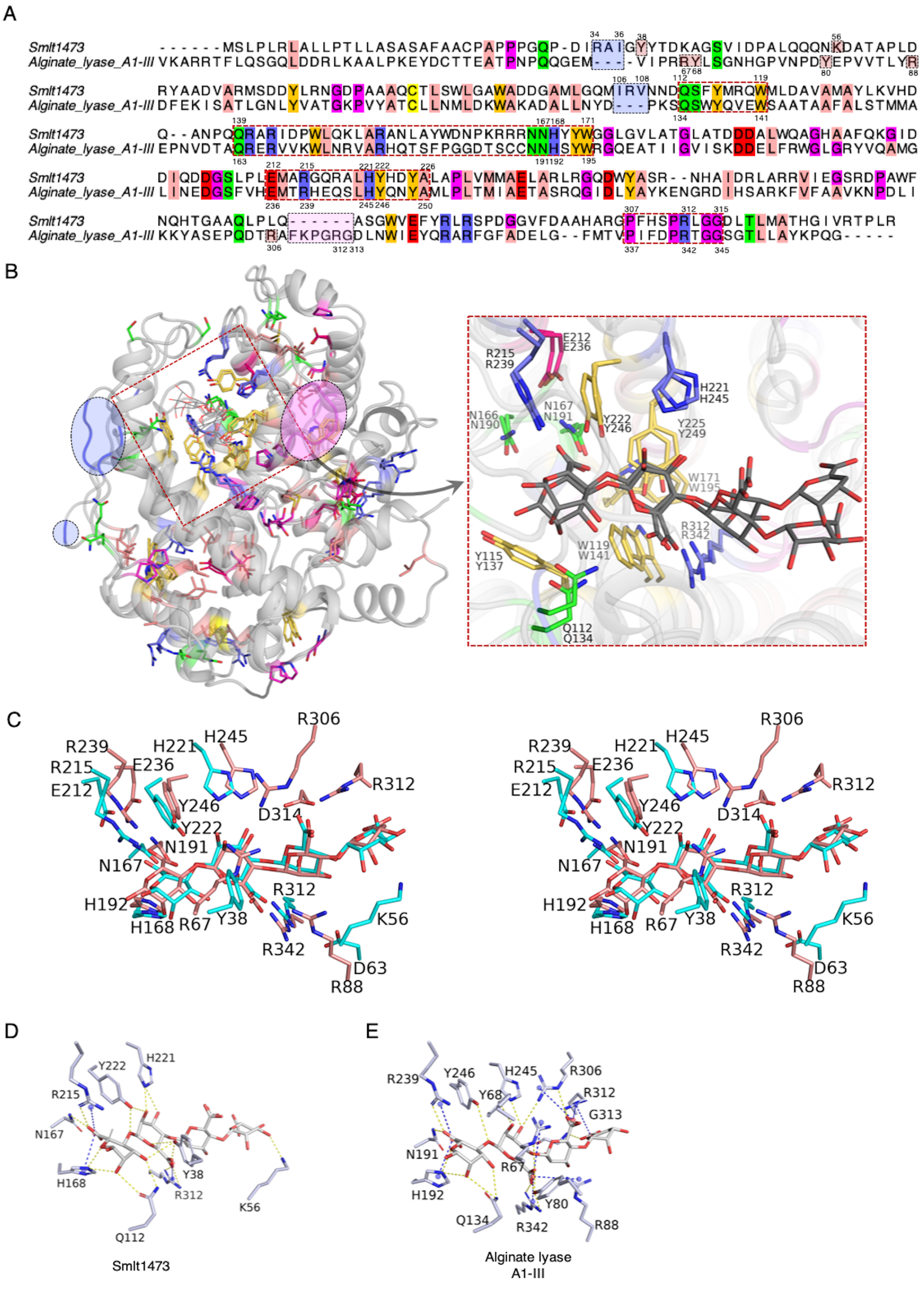

**FIGURE S11. Sequence-Structure comparison of PL-5 family of polyManA lyases**. **(A)** Structure guided sequence alignment of Smlt1473 (multi-functional), and Alginate lyase A1-III (polyManA specific) the conserved residues are highlighted by amino acid color code as discussed in Figure S4. The active site is highlighted under a red dashed box, the Smlt1473's insertion at N-terminal are highlighted under light blue box, and Alginate lyase A1-III's C-terminal insertion is highlighted under light purple box. **(B)** Structural alignment of Smlt1473, and Alginate lyase A1-III is displayed as cartoon, the conserved amino acids are displayed as sticks which are color coded according to type of amino acids described in A, the region of sequence insertion cartoon model are colored and labelled as described in A. The conserved active site region based on sequence alignment are marked under red dashed box, which has been zoomed to show the conserved amino acid residues. **(C)** Stereographic image of titrable amino acids around the structurally aligned substrate complexed with Smlt1473 (cyan stick), and Alginate lyase A1-III (salmon stick). **(D-E)** Enzyme-Substrate interaction derived from modelled (catalytic base is computationally mutated back to wildtype) and relaxed (to remove steric clashes) substrate (ManA) complexed structure of selected PL-5 enzymes.

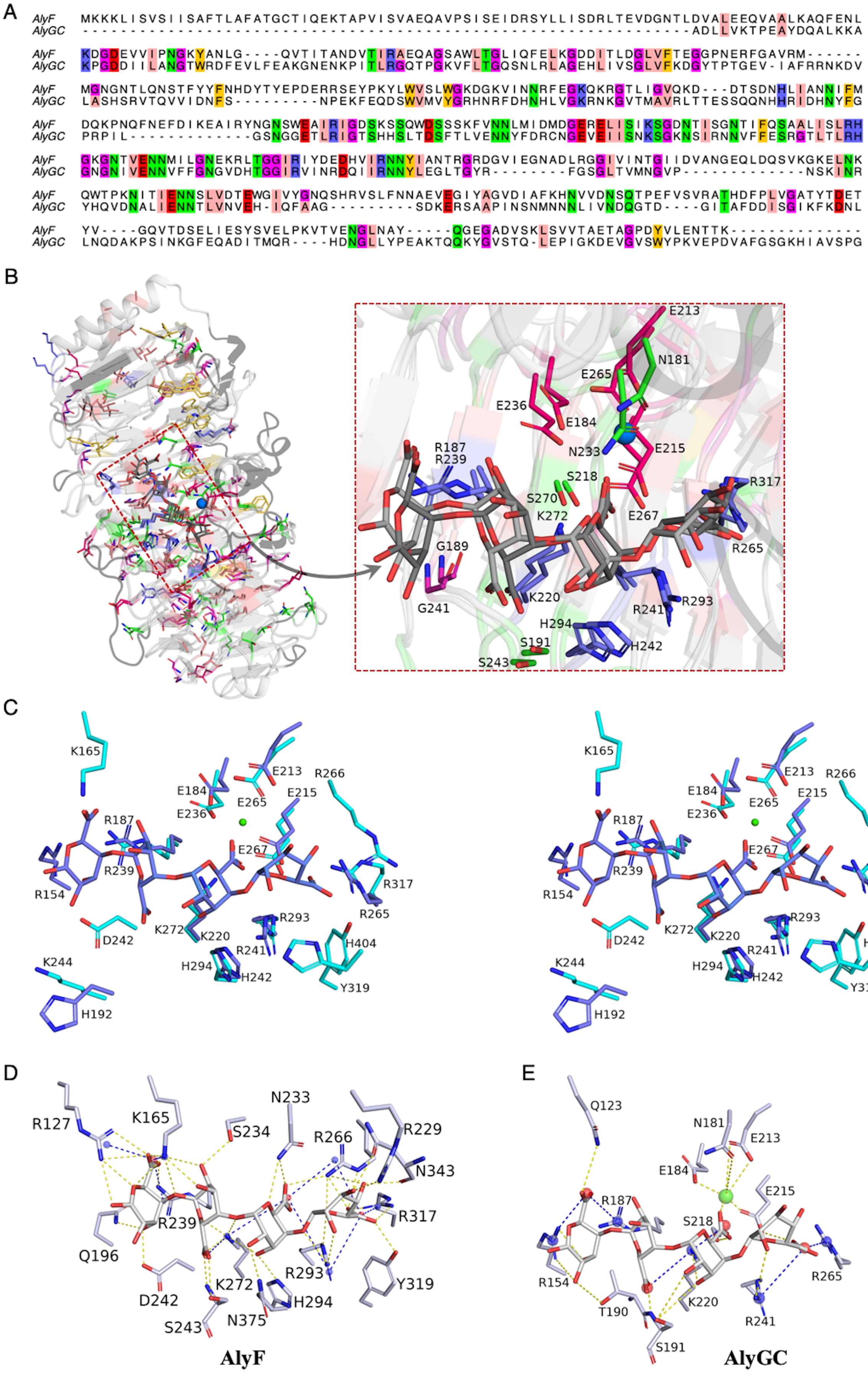
A
AlyF MKKKLISVSIISAFTLAFATGCTIQEKTAPVISVAEQAVPSISEIDRSYLLISDRLTEVDGNTLDVALEEQVAALKAQFENL
AlyGC ---------------------------------------------------------------ADLLVKTPEAYDQALKKA
AlyF KDGDEVVIPNGKYANLG----QVTITANDVTIRAEQAGSAWLTGLIQFELKGDDITLDGLVFTEGGPNERFGAVRM------
AlyGC KPGDDIILANGTWRDFEVLFEAKGNENKPITLRGQTPGKVFLTGQSNLRLAGEHLIVSGLVFKDGYTPTGEV-IAFRRNKDV
AlyF MGNGNTLQNSTFYYFNHDYTYEPDERRSEYPKYLWVSLWGKDGKVINNRFEGKQKRGTLIGVQKD---DTSDNHLIANNIFM
AlyGC LASHSRVTQVVIDNFS---------NPEKFEQDSWVMVYGRHNRFDHNHLVGKRNKGVTMAVRLTTESSQQNHHRIDHNYFG
AlyF DQKPNQFNEFDIKEAIRYNGNSWEAIRIGDSKSSQWDSSSKFVNNLMIDMDGERELISIKSGDNTISGNTIFQSAALISLRH
AlyGC PRPIL-------------GSNGGETLRIGTSHHSLTDSFTLVENNYFDRCNGEVEIISNKSGKNSIRNNVFFESRGTLTLRH
AlyF GKGNTVENNMILGNEKRLTGGIRIYDEDHVIRNNYIANTRGRDGVIEGNADLRGGIVINTGIIDVANGEQLDQSVKGKELNK
AlyGC GNGNIVENNVFFGNGVDHTGGIRVINRDQIIRNNYLEGLTGYR--------FGSGLTVMNGVP-------------NSKINR
AlyF QWTPKNITIENNSLVDTEWGIVYGNQSHRVSLFNNAEVEGIYAGVDIAFKHNVVDNSQTPEFVSVRATHDFPLVGATYTDET
AlyGC YHQVDNALIENNTLVNVEH-IQFAAG---------SDKERSAAPINSNMNNNLIVNDQGTD----GITAFDDISGIKFKDNL
AlyF YV----GQVTDSELIESYSVELPKVTVENGLNAY-----QGEGADVSKLSVVTAETAGPDYVLENTTK---------------
AlyGC LNQDAKPSINKGFEQADITMQR----HDNGLLYPEAKTQQKYGVSTQ-LEPIGKDEVGVSWYPKVEPDVAFGSGKHIAVSPG
B
E213
N181
E265
E236
E184
R187
R239
N233
E215
S218
R317
S270
K272
E267
G189
R265
K220
G241
R241
R293
H294
S191
H242
S243
C
K165
E213
R266
E184
E215
E236
E265
R187
E267
R239
R317
R154
R265
K272
K220
R293
D242
R241
H404
K244
H294
H242
Y319
H192
D
R127
K165
N233
S234
R266
R229
N343
R239
R317
Q196
R293
Y319
K272
H294
D242
N375
S243
AlyF
E
Q123
N181
E213
E184
E215
R187
S218
R154
R265
K220
T190
R241
S191
AlyGC

**FIGURE S12. Sequence-Structure comparison of PL-6 family of polyGulA lyases**. **(A)** Structure guided sequence alignment of AlyF ($Ca^{2+}$ ion independent, polyGulA specific PL), and AlyGC (bi-functional alginate lyase) the conserved residues are highlighted by amino acid color code as discussed in Figure S4. **(B)** Structural alignment of AlyF, and AlyGC is displayed as cartoon, the conserved amino acids are displayed as sticks which are color coded according to type of amino acids described in A, the region of sequence insertion cartoon model are colored black. The conserved active site region based on sequence alignment are marked under red dashed box, which has been zoomed to show the conserved amino acid residues. **(C)** Stereographic image of titrable amino acids around the structurally aligned substrate complexed with AlyF (cyan stick), and AlyGC (blue stick). **(D-E)** Enzyme-Substrate interaction derived from modelled (catalytic base is computationally mutated back to wildtype) and relaxed (to remove steric clashes) substrate (ManA) complexed structure of selected PL-6 enzymes.

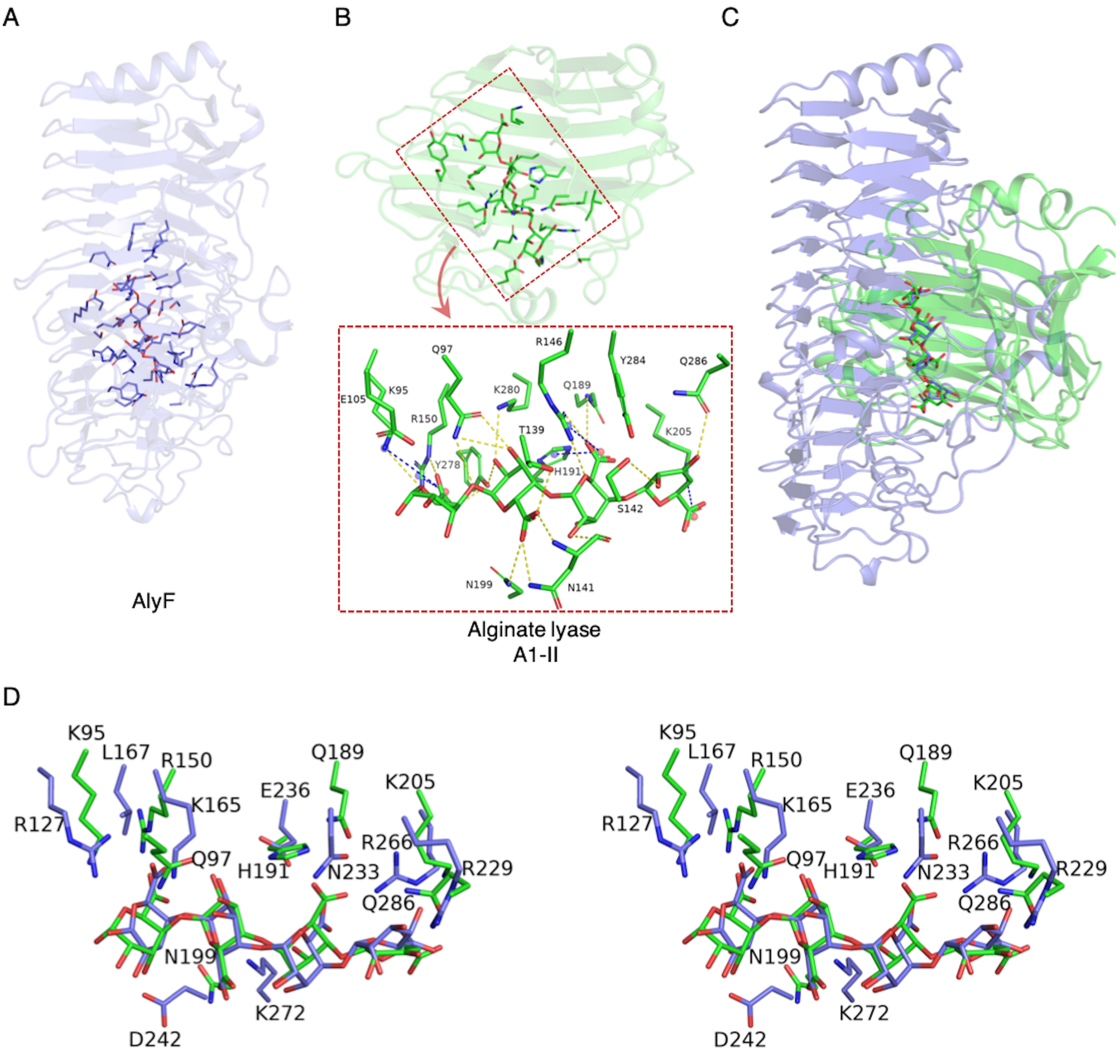

**FIGURE S13. Structural comparison of PL7 and PL-6 family of PLs cleaving polyGulA. (A)** Structural description of AlyF as cartoon model with active site depicted as stick, **(B)** Structural description of Alginate lyase A1-II as cartoon model with active site depicted as stick and zoomed inset (red dashed box) depict the enzyme-substrate interaction. **(C)** Substrate coordinate based structural alignment of AlyF (blue), and Alginate lyase A1-II (green) shown as cartoon model, **(D)** stereo image of the active site's titrable amino acid around the structurally aligned complexed GulA from AlyF and Alginate lyase A1-II.
